# Temporally overlapping mechanisms diversify clonal B cell responses *in vivo*

**DOI:** 10.1101/2024.12.17.628863

**Authors:** Oliver P. Skinner, Saba Asad, Marcela L. Moreira, Hyun Jae Lee, Cameron G. Williams, Zheng Ruan, Jorene Lim, Ashlyn S. Kerr, Shihan Li, Chen Zhu, Wang Jin, Thiago M. Steiner, Takahiro Asatsuma, Brooke J Wanrooy, Zihan Liu, Marcus Z.W. Tong, Megan S.F. Soon, Jessica A. Engel, Kalyan Shobhana, Adam P. Uldrich, David S. Khoury, Zewen Kelvin Tuong, Hamish W. King, Ashraful Haque

## Abstract

Naive B cells amplify and diversify their responses when activated by cognate antigen, via Myc-dependent clonal expansion, immunoglobulin class switch recombination (CSR), phenotypic variation, and somatic hypermutation (SHM). Whether these mechanisms act combinatorially in vivo to diversify clonal responses to a single pathogen remains unclear. Since diversity in the antigenic targets, functional classes, and production kinetics of parasite-specific antibodies influences immunity to malaria, we test here whether individual B cell clones diversify over time during Plasmodium infection and treatment. During the first week of infection, amid widespread Type I Interferon (IFN)-mediated bystander activation, CSR initiates soon after Myc up-regulation, and overlaps partially with clonal expansion, resulting in isotype variegation amongst clones. During the second week of infection, expanded clones that seed germinal centres (GC) bifurcate into extra-follicular plasmablasts, exhibit isotype variegation, and initiate SHM, revealing substantial intra-clonal diversification. Over the following month, GC clones exhibit SHM at approximately four mutations per week, with IgG mutational diversity and IgM^+^ cells also preserved in GCs over time. Anti-malarial intervention does not impede SHM, but instead exerts quantitative limits on GC size, plasma cell emergence, circulating IgG levels, and protection against re-infection. Finally, contemporaneous B cell development relocates from bone marrow to spleen during infection and treatment. Thus, multiple temporally overlapping mechanisms combine in vivo to amplify, diversify, and safeguard humoral immune responses. We present this data as a temporal, multi-parameter atlas of B cell differentiation in vivo: https://bcell-dynamics.science.unimelb.edu.au

**Graphical abstract:** 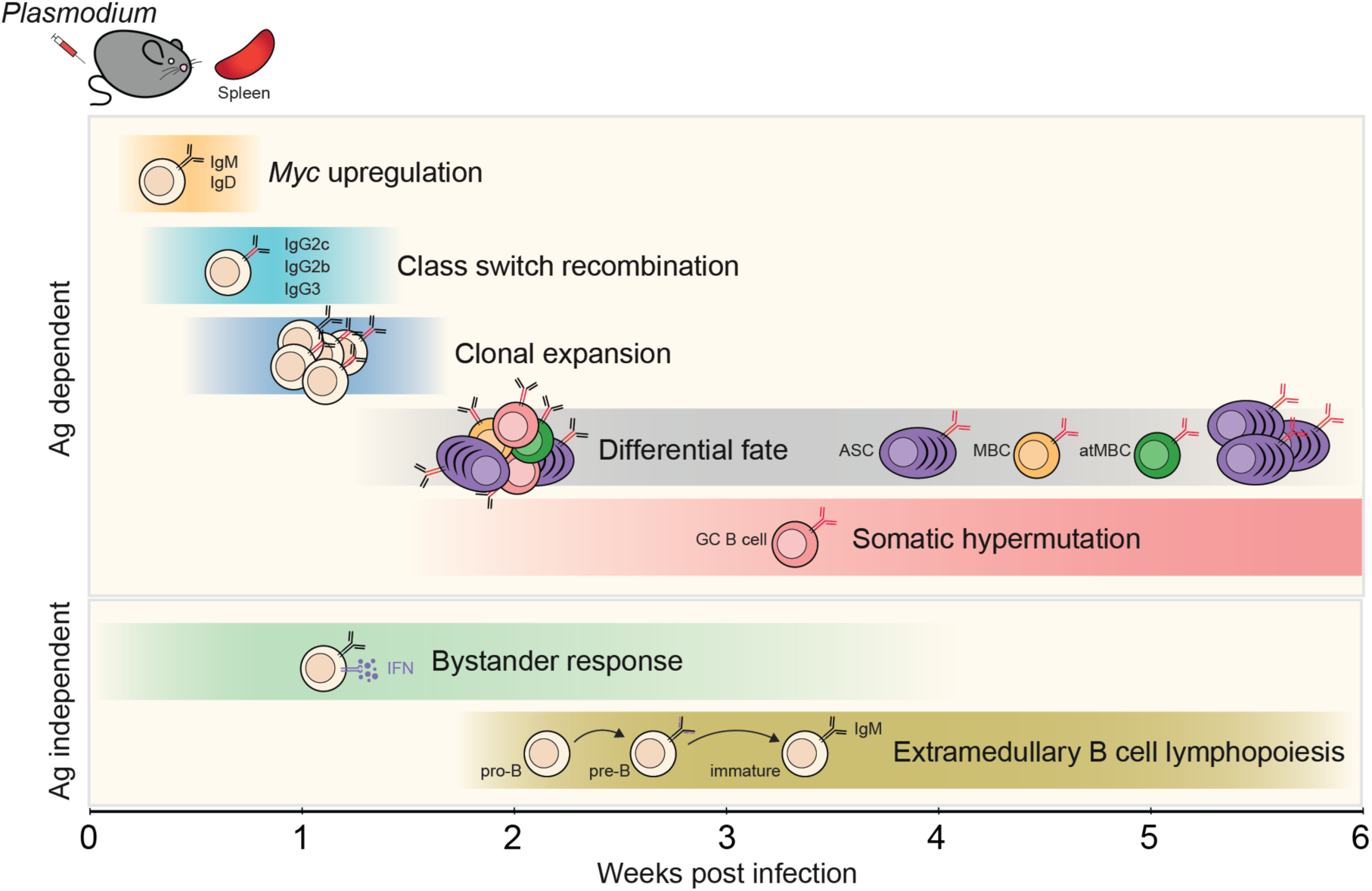

**Highlights:**

- Partial temporal overlap of CSR with clonal expansion leads to isotype variegation in clones.
- Clones seeding GCs bifurcate into plasmablasts and exhibit isotype variegation.
- GC B cells accrue ∼4 mutations/week, a rate unaffected by anti-malarials.
- *Plasmodium* infection triggers antigen-independent Type I IFN-mediated bystander activation.
- B cell development is preserved in malaria by shifting from bone marrow to spleen.

## Introduction

Antigen receptor engagement on naive B cells initiates not only clonal expansion, but also a range of phenotypic changes and genetic alterations that increases diversity beyond that offered by pre-existing BCRs alone. Phenotypes adopted include short-lived antibody-secreting plasmablasts, memory B cells, atypical B cells, germinal centre B cells and plasma cells, each with distinct functions within the overall humoral immune response. Further diversification proceeds via class switch recombination (CSR), which replaces the constant portion of BCRs with one of a number of functionally varied isotypes (*e.g.* IgG1, IgG2, IgG3, IgG4, IgA1, IgA2 and IgE in humans)^1,2^. In addition, B cells entering germinal centres (GC) can further diversify their BCRs via somatic hypermutation (SHM)^3-6^, leading to affinity maturation of resulting antibodies^7^.

B cell diversification during infection can either promote or impair humoral immunity, as evidenced during blood-stage *Plasmodium* infection, in which plasmablast responses can limit immunity to malaria^8^, while increased breadth of antibody specifities^9,10^ and effector functions^11^ is associated with greater protection against symptoms. Understanding when diversification mechanisms operate and whether they interact *in vivo* may offer opportunities for tuning immunological responses. In addition, although anti-malarial drugs have been instrumental in reducing the morbidity and mortality associated with malaria^12^, their impact on long-term immunity remains unclear^13-18^.

Experimental vaccination with single antigens has revealed several *in vivo* aspects of CSR and phenotypic diversification in single naive B cells. Firstly, CSR can occur prior to GC entry ^19-21^. Secondly, in some cases^22^, but not others^23^, CSR occurs independently of phenotypic changes. Thirdly, the degree of phenotypic diversification by expanded clones is controlled by the BCR itself, and is further regulated by apoptosis during clonal expansion^22^. CSR is largely thought to act orthogonally to processes determining phenotypic fate, relying on stochastic expression of activation-induced cytidine deaminase (AID) and germ-line transcripts (GLT)^24^. Viral infection studies have revealed SHM dynamics within the GC^25^, showing that mutations accrue and then diminish due to ongoing invasion by naive B cells^25,26^. Recent studies of steady-state human tonsils identified transient, intermediate states of B cell activation^27,28^. Although these observations provide vignettes of B cell clonal diversification, temporal relationships of, and interactions between these mechanisms remain to be determined within a single *in vivo* model.

Here, we employ scRNAseq, BCRseq, and spatial transcriptomics to map over time the diversification mechanisms that operate in polyclonal B cells *in vivo,* using a model of blood-stage infection with *Plasmodium* parasites, an antigenically complex protozoan^29,30^, which continues to threaten global human health^31^.

## Results

### Plasmablast and GC B cell differentiation temporally overlaps with Type I IFN-mediated bystander activation during experimental malaria

To assess polyclonal B cell diversification over time, we employed blood-stage *Plasmodium chabaudi chabaudi* (*Pc*AS) infection of C57BL/6J mice. This model is characterised by peak parasitemia ∼7-9 days post-infection (dpi), followed by CD4^+^ T-cell dependent, antibody-mediated, parasite elimination over the following 60-90 days^32,33^. We performed paired scRNAseq and BCRseq of splenic polyclonal B cells (CD19^+^ B220^int-hi^) prior to and 4-, 7-, 10-and 14 dpi, reasoning this would capture early activation events and progressive emergence of plasmablasts and GC B cells (Fig. 1A). From 20 mice, 40,623 transcriptomes passed quality-control: day 0 (2,322 cells, n=1); day 4 (9,079 cells, n=4), day 7 (4,337 cells, n=5), day 10 (11,407 cells, n=5) and day 14 (13,478 cells, n=5). Dimensionality reduction, unbiased clustering, and canonical marker analysis revealed multiple subsets emerging over time (Extended Data Fig. 1A-C), in particular plasmablasts and GC B cells evident at days 10 and 14 dpi, displaying kinetics consistent with previous flow cytometric studies^32,33^ (Fig. 1B). This suggested the data could be employed for examining B cell diversification mechanisms.

**Figure 1:**
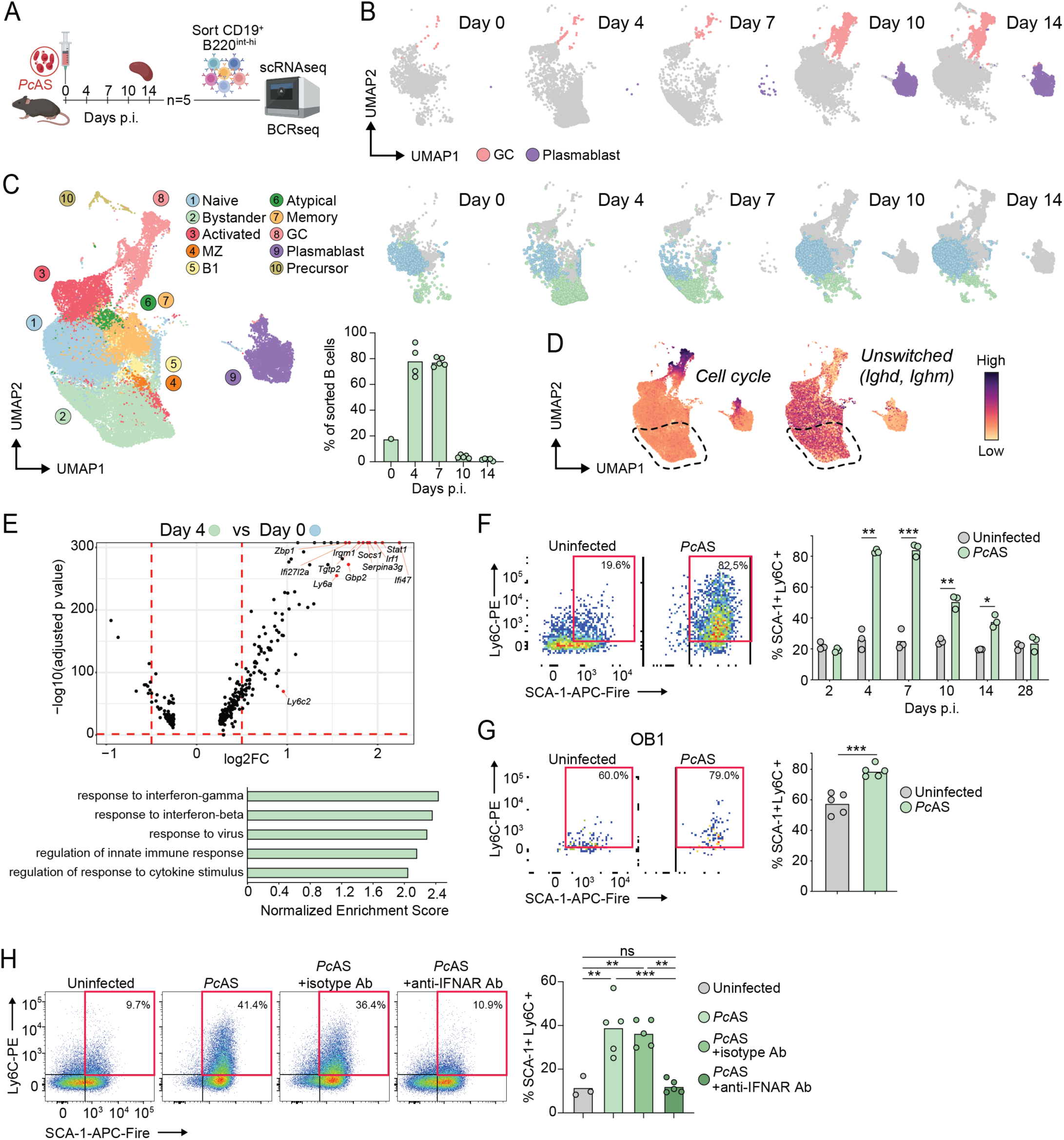
Plasmablast and GC B cell differentiation temporally overlaps Type I IFN-mediated bystander activation during experimental malaria. (**A**) Experimental plan to study B cell differentiation in *Pc*AS-infected mice. Splenic CD19^+^ B220^int-hi^ B cells were isolated at various timepoints post-infection and underwent droplet-based, paired single cell RNA and BCR sequencing. (**B**) High-dimensional clustering UMAP of GC B cells (pink) and plasmablasts (purple). (**C**) UMAP plot of combined B cell populations across all (left) or specific timepoints (right) with the two follicular B cell subsets; naive (blue) and bystander (green) and all other B cells (grey). Frequency of bystander B cells at each timepoint are indicated by the bar graph (middle). (**D**) Gene signature feature plots of cell cycle or unswitched signatures (dashed lines highlight bystanders). (**E**) Volcano plot highlighting the top differentially upregulated genes in red expressed in follicular (day 4) compared to naïve (day 0) clusters. Gene set enrichment analysis (GSEA) highlights the top five significantly enriched biological processes gene ontology (GO) terms. (**F**) Representative pseudocolour density plots of SCA-1 and Ly6C expression ("bystander markers") in IgD^+^ B cells at day 4 post-*Pc*AS infection (left) and bar graph of SCA-1^+^ Ly6C^+^ cell frequencies amongst IgD^+^ B cells over time (right). (**G**) Representative pseudocolour density plots (left) and bar graph (right) showing proportion of SCA-1 and Ly6C expression on CD86^lo^ OB1 adoptively transferred B cells at 4 days post-*Pc*AS infection. (**H**) Representative pseudocolour density plots (left) and bar graph (right) of SCA1 and Ly6C-expressing IgD^hi^ CD19^+^ B220^+^ B cell frequencies at day 4 post-*Pc*AS infection in the presence of continual anti-IFNAR antibody blockade or isotype control. Data from bar graphs in C-H represent the mean, where F-G were compared using Welch’s *t*-test and H by one-way ANOVA with Tukey’s multiple comparisons test. All experiments are representative of two independent experiments of *n =* 3-5 per group. \**P* <0.05, \*\**P* <0.01, \*\*\**P* <0.001. ns, not significant.

In addition to expected plasmablast and GC B cell differentiation we noted increased frequency at 4 and 7 dpi of a cluster of unswitched, non-proliferating B cells (Fig. 1C-D), defined by up-regulation of interferon-associated genes, including *Stat1*, *Socs1, Irf1* and *Irgm1* (Fig. 1E). *Ly6a* (encoding SCA-1) and *Ly6c2* (encoding Ly6C), were also upregulated (Extended Data Fig. 2A). This suggested B cells lacking specificity for *Plasmodium* antigens had responded in a bystander, interferon-mediated fashion during infection. Kinetics of SCA-1 and Ly6C expression were confirmed at protein level on IgD^+^ cells by 4 dpi, which was sustained until day 14 (Fig. 1F). SCA-1/Ly6C up-regulation was less pronounced in CD4-depleted mice (Extended Data Fig. 2B)^32^, suggesting partial T-cell-dependence for this phenomenon. We confirmed bystander activation in ovalbumin-specific OB1 cells after transfer and infection with non-ovalbumin expressing *Pc*AS (Fig. 1G; Extended Data Fig. 2C). Finally, SCA-1/Ly6C upregulation by IgD^+^ cells was abrogated by IFNAR1-blockade (Fig. 1H), while B cell-intrinsic genetic deficiency of Toll-like receptor adaptor, *Myd88,* had no effect (Extended Data Fig. 2D). Thus, plasmablast and GC B cell differentiation had occurred amid Type I IFN-mediated bystander B cell activation in the spleen during experimental malaria.

### CSR occurring soon after *Myc*-upregulation overlaps with clonal expansion

Given plasmablasts and GC B cells had been detected within the scRNAseq data by 10-14 dpi (Fig. 1B), we next searched for evidence of early activation, clonal expansion and progressive phenotypic change. We observed a transient increase at 4 dpi of cells marked by upregulation of *Myc, Nfkbid*, *Cd83, Cd69* and *Jun*, consistent with early activation^27,28,34-36^ (Fig. 2A). Although we observed no cell-states consistent with intermediate or progressive differentiation towards plasmablasts, we did note early activated cells positioned adjacent to emerging GC B cells (Fig. 2A). Closer examination revealed that early-activated cells, in addition to strong *Myc-*expression had up-regulated CSR-related genes^27^ though not cell-cycling genes (Fig. 2B). Inference of progressive, overlapping, transitions from *Myc-*upregulation to CSR and clonal expansion was supported by Bayesian Gaussian Process Latent Variable Modelling (BGPLVM) and pseudo-temporal ordering (Fig. 2B & C), an approach we previously used for inferring progressive change in CD4^+^ T cells^37-39^. Given only a subset of cells within the early activated cluster had upregulated *Myc* and CSR-related genes, we re-clustered these to further test for overlap (Extended Data Fig. 3A), and to search for novel genes associated with early activation. After unsupervised clustering, we noted Cluster 5 exhibited the highest *Myc-*upregulation (Extended Data Fig 3B), amongst other genes (Extended Data Fig. 3C, Supplementary Table 1), while Cluster 8 displayed moderate *Myc-*upregulation, the highest expression of CSR-related genes, and emerging upregulation of cell-cycling genes, within activated B cells (Extended Data Fig. 3B), consistent with temporal overlap.

**Figure 2:**
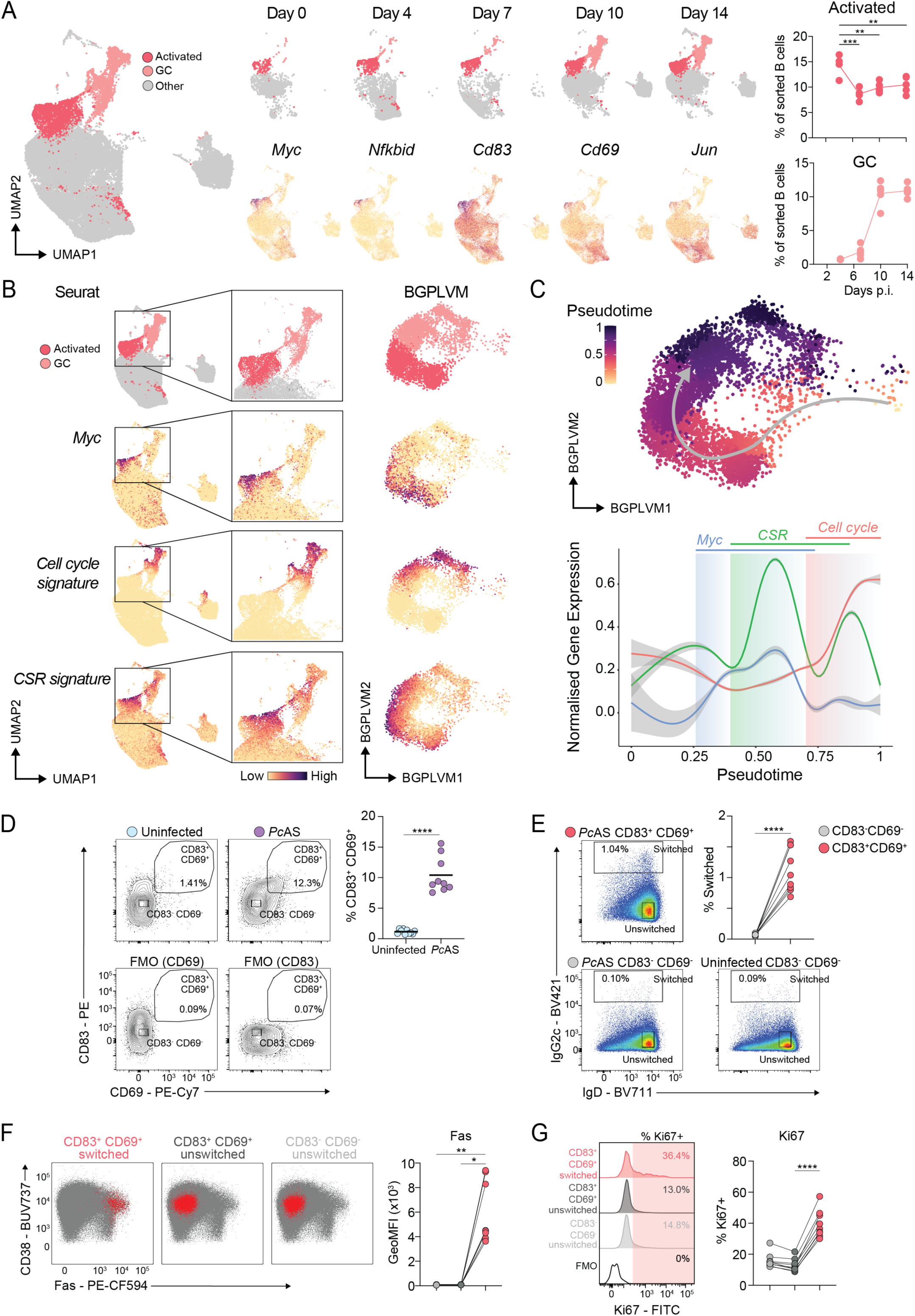
CSR occurring soon after *Myc*-upregulation overlaps with clonal expansion. (**A**) Annotated B cell UMAP (left) of activated (indicated by activation marker gene expression: *Myc, Nfkbid, Cd83, Cd69* and *Jun*) and GC B cells after high-dimensional reduction. Clustering is additionally split by each timepoint (middle) with quantitation of replicate mice across time for activated and GC B cells (right). (**B**) Feature plots of *Myc* expression, Class Switch Recombination (CSR) and cell cycle gene signatures across Seurat and Bayesian Gaussian Process Latent Variable Model (BGPLVM) embedding methods. (**C**) Pseudotime trajectory analysis of BGPLVM object using GPFates, including *Myc*, CSR signature and cell cycle signature-normalised expression along pseudotime. (**D**) Representative contour plots (left) and scatter plot (right) of CD83^+^ CD69^+^ activated B cell frequencies as gated on CD19^+^ B220^+^, non-GC B cells and non-PBs, in uninfected and *Pc*AS (4 dpi) from total B cells. Activated B cell gates were drawn based on CD69^-^ and CD83^-^FMOs. (**E**) Representative pseudocolour density plots (left) of IgG2c^+^ switched CD83^+^ CD69^+^ (activated) and CD83^-^ CD69^-^ B cells frequencies (right) from *Pc*AS-infected samples. (**F**) Representative dot plots (left) of Fas expression in selected cell populations (highlighted in red) and line graph (right) of geometric mean fluorescence intensity (GeoMFI) of Fas in populations of CD83^+^ CD69^+^ switched, CD83^+^ CD69^+^ unswitched, and CD83^-^ CD69^-^ unswitched B cells. (**G**) Representative histograms (left) and line graph (right) of % Ki67^+^ cells in populations of CD83^+^ CD69^+^ switched, CD83^+^ CD69^+^ unswitched and CD83^-^ CD69^-^ unswitched B cells. Ki67^+^ gate drawn based on Ki67^-^ FMO. Lines connect the mean value in panel a at each timepoint and analysed by one-way ANOVA with Tukey’s multiple comparisons test. Data in panel D-G is pooled from two independent experiments of *n* = 9-10 per group. Data in panel D represents the mean and analysed using Welch’s *t*-test. Data in panel E analysed using a paired *t*-test. Data in panels F-G analysed using Friedman test with Dunn’s multiple comparisons test. \**P* <0.05, \*\**P* <0.01, \*\*\**P* <0.001, \*\*\*\**P* <0.0001.

Finally, we tested for early overlap of CSR and clonal expansion by flow cytometric assessment of splenocytes at 4 dpi. First, co-expression of CD83 and CD69 was employed to enrich for polyclonal B cells recently activated by cognate antigen (Fig. 2D). Next, compared to CD83^-^ CD69^-^ B cells in either infected or naïve control mice, ∼1% of recently activated CD83^+^ CD69^+^ cells had switched to IgG2c (Fig. 2E). Of note, the majority of these co-expressed IgD, which given the relatively low turnover of surface protein IgD^40^, suggested these cells had recently switched. Furthermore, these cells expressed Fas, similar to a report in which Fas^+^ B1-8^hi^ cells underwent CSR prior to GC seeding^21^ (Fig. 2F). Crucially, IgG2c^+^ CD83^+^ CD69^+^ recently-switched B cells exhibited elevated levels of the proliferative marker, Ki67 relative to unswitched comparator populations (Fig. 2G), consistent with overlap of CSR and clonal expansion. Together, our data support a model in which CSR initiates soon after *Myc* upregulation, and importantly, partially overlaps with clonal expansion.

### Activated B cell clones seeding GCs bifurcate into plasmablasts and exhibit isotype variegation

Partial temporal overlap of CSR and clonal expansion suggested a proportion of activated B cell clones, having undergone at least one cell division prior to CSR, could exhibit Ig isotype variegation (since our model elicits IgG2b, IgG2c, IgG1, IgG3, and IgM^32,33,41^). Furthermore, given subsequent differentiation into plasmablasts and GC B cells (Fig. 1B), we hypothesized single naïve B cells could elicit diverse progeny exhibiting both multiple fates and multiple Ig-isotypes.

To test for such clonal diversification, we first searched for expanded clones in our initial 0-14 dpi scRNAseq/BCRseq dataset. Cells were defined as clonal when sharing identical *V* and *J* genes, identical junctional CDR3 amino acid length, and >85% amino acid CDR3 sequence identity as previously described^42,43^. Amongst ten mice across 10 and 14 dpi, we detected 356 expanded clones harbouring 2-8 cells (Fig. 3A), the majority of which (297 clones) consisted of two cells (hereafter referred to as “twins”), which were shared relatively uniformly across all mice (Fig. 3B). A repeat scRNAseq/BCRseq experiment (6 mice across 10 & 14 dpi) supplemented the dataset, providing a total of 721 expanded clones, of which 592 were twins (Fig. 3B & C). Thus, despite shallow sampling of splenic B cell compartments in individual mice, our datasets nevertheless permitted testing for clonal diversification.

**Figure 3:**
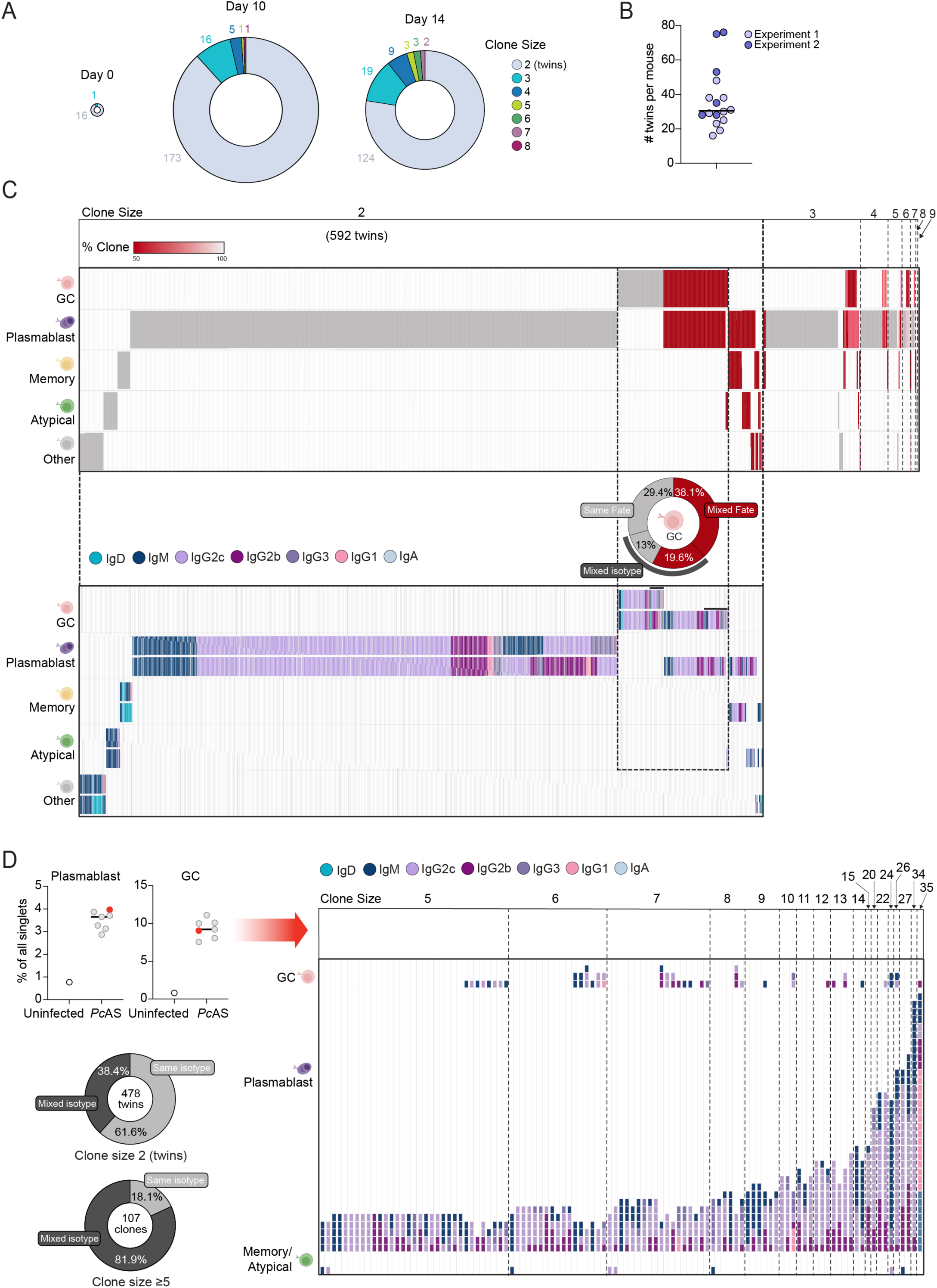
Activated B cell clones seeding GCs bifurcate into plasmablasts and exhibit isotype variegation. (**A**) Donut plots highlighting clone size distribution of expanded B cell clones detected at 0, 10 and 14 dpi. Size of donut plot is proportional to number of expanded clones at each timepoint. (**B**) scatter plot of the number of twin clones detected per mouse at 10 and 14 dpi from two experiments. (**C**) Clonal fates ordered by clonal size of plasmablast, GC, memory, atypical and other B cell subsets at 10 and 14 dpi. Each row represents cell fate, and each column represents the frequency of cells per clone. Clones are split by same fate (grey) and mixed fate (red). Donut plot represents quantification of same (grey) and mixed fates (red) in GC-containing clones of size 2. Clones of size 2 (of which there were 592 twins in total) were further analysed for their isotype in each cell, with donut chart of same and mixed isotypes (denoted with thick black line) for each clone with same (grey) and mixed fate (red) outcomes. (**D**) A single mouse spleen which had representative proportion of plasmablasts and GC B cells at 14 dpi (denoted in red), underwent paired single cell RNA and BCR sequencing (left). Plot on the right highlights the 107 expanded clones (≥ 5). Each column represents a clone, and each row depicts the cell fate within each clone, coloured by isotype. Donut charts highlight the proportion of clones (twins or clones greater than 5) with same and mixed isotypes.

Firstly, we determined across 10 and 14 dpi whether individual clones could differentiate into both GC B cells and plasmablasts (Fig. 3C). The majority (567 out of 721) of expanded clones gave rise to plasmablasts only, consistent with these short-lived antibody secreting cells dominating at early time-points, although we noted ∼50% reduced efficacy in BCR-detection amongst GC B cells compared to plasmablasts due to lower BCR gene expression levels. Nevertheless, across all 461 plasmablast twins, 119 (25.8%) of these exhibited isotype variegation, consistent with partial temporal overlap of CSR and clonal expansion (Fig. 3C).

Amongst 592 twins, 92 gave rise to at least one GC B cell, with 53 (57.6%) bifurcating to give rise to a plasmablast (Fig. 3C) – the frequency of fate bifurcation rose to 100% for the small number (19 clones) of GC-containing clones with >2 family members (Fig. 3C). Furthermore, of GC-containing twins 19.6% exhibited both isotype variegation and fate bifurcation (Fig. 3C), consistent with our original hypothesis.

Finally, to examine clonal diversification in larger clone sizes, we sampled more deeply plasmablasts and GC B cells flow cytometrically sorted from a single representative spleen (n=5) at 14 dpi (Fig. 3D, Extended Data Fig. 4A). From 19,482 post-QC B cell transcriptomes from a single mouse, 9,350 plasmablasts, and 8,744 GC B cells were detected in the paired scRNAseq/BCRseq data (with minor populations of memory/atypical cells and spike-in naive cells) (Extended Data Fig. 4A). Amongst these, we detected 478 twins, 38.4% of which exhibited isotype variegation (Fig. 3D), further supporting temporal overlap for CSR and clonal expansion. Of 107 clones comprised of 5 cells or more, and similar to experiments above (Fig. 3C), 100% of GC-containing clones (34 clones) had also given rise to plasmablasts (Fig. 3D) (which harboured as few mutations as naïve B cell controls, consistent being extrafollicular, not GC-derived) (Extended Data Fig. 4B). Furthermore, isotype variegation was evident in most (81.9%) clones comprised of 5 cells or more (Fig. 3D). Taken together, our data suggested it was commonplace during the first two weeks of infection for antigen-activated B cells, particularly GC-seeding clones to have exhibited both phenotypic and Ig-isotype diversification.

### GC B cells progressively diversify via a constant SHM rate and retention of isotype diversity

Having observed phenotypic and isotype diversification amongst GC-seeding clones, we next sought to determine the extent to which SHM further diversified B cells in our model of persisting infection. Furthermore, given evidence of reduced affinity maturation and disrupted splenic GC structures in malaria-endemic regions^44,45^, we also sought to determine if anti-malarial drug intervention altered GC B cell biology during *Plasmodium* infection. Thus, we generated an extended scRNAseq/BCRseq time course until 42 dpi, which included a parallel arm of anti-malarial drug intervention from 7 dpi (Fig. 4A, Extended Data Fig. 5A).

**Figure 4:**
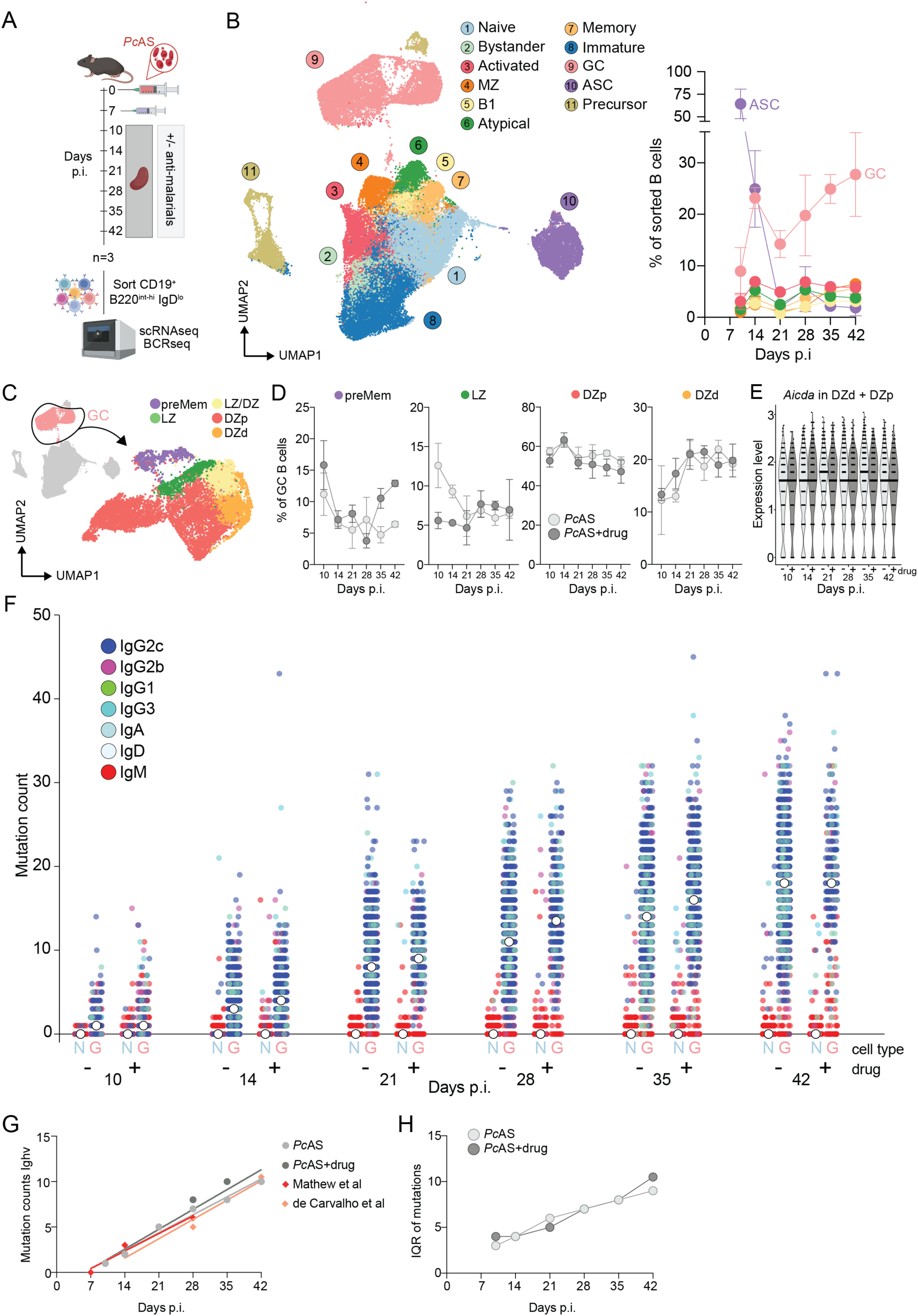
GC B cells progressively diversify via a constant SHM rate and retention of isotype diversity. (**A**) Experimental plan to study B cell differentiation in *Pc*AS-infected mice, with or without anti-malarial drug treatment. CD19^+^ B220^int-hi^ IgD^lo^ splenic B cells isolated at various timepoints post-infection underwent droplet-based, paired single cell RNA and BCR sequencing. (**B**) Annotated combined UMAP of B cell subsets from all mice, treatments and timepoints after high-dimensional reduction and clustering (left), and line graph (right) of selected, annotated B cell frequencies over the course of *Pc*AS infection without anti-malarial drug treatment. (**C**) Sub-clustering of the GC cluster identifies preMem, LZ, LZ/DZ, DZp and DZd B cells. (**D**) Line graphs of GC B cell subset frequencies over time, with or without drug treatment. (**E**) Violin plot of the median *Aicda* expression shown over time, with or without anti-malarial treatment in DZd and DZp GC B cells. (**F**) Scatter plot of somatic hypermutation rates of naïve (N) and GC (G) B cell BCRs and isotype usage between drug or non-drug-treated mice over time. Median number of mutations for each treatment and timepoint indicated in white. (**G**) Line graph of interquartile range (IQR) of GC B cell mutations at each timepoint. (**H**) Line graph of median number of mutations in the *Ighv* region of the BCR for each condition in comparison to clonal data from lymph node GC B cells after influenza infection from Mathew et al. and de Carvalho et al. Data in B and D are shown as mean ± SD of *n* = 3 per group.

To focus assessment on B cells activated via cognate antigen, we FACS-sorted IgD^lo^ B cells (CD19^+^ B220^int-hi^), spiking back in small numbers of IgD^hi^ naive cells at each timepoint to assist in batch effect removal. We isolated 72,359 B cells across two treatment arms (*Pc*AS: 38,651 cells, *Pc*AS+drug: 27,452 cells, n=3 mice per timepoint) from 42 mice: IgD^hi^ spike-in from un-infected mice (6,256 cells, n=7); 10 dpi (9,250 cells), 14 dpi (10,338 cells), 21 dpi (15,766 cells), 28 dpi (12,251 cells), 35 dpi (15,502 cells) and 42 dpi (9,252 cells). Within the combined scRNAseq data (after confirmation of minimal batch effects), we inferred 11 distinct cell types including transient ASCs and increasing GC B cell frequencies (Fig. 4B). Most cell-types showed little change in frequency as a result of anti-malarial treatment, except for GC B cells which dropped by 36.2%, and MZ B cells and B1 cells, which increased by 148% and 75%, respectively (Extended Data Fig. 5B). Flow cytometric assessment confirmed a reduction in GC B cell numbers within the spleen by 42 dpi following drug treatment, with similar trends for ASCs, atypical and memory B cells (Extended Data Fig. 5C) as observed in the scRNAseq data (Extended Data Fig. 6A-B).

We next searched for qualitative changes in GC B cells over time and after anti-malarial treatment within the extended scRNAseq time-course. Phenotypic characterisation of GC B cell subsets using canonical gene markers and gene signatures^46^, identified putative Light Zone (LZ), Dark Zone proliferating (DZp) and differentiating (DZd) cells, as well as mixed LZ/DZ cells and *Ccr6^+^* memory B cell precursors (preMems) (Fig. 4C, Extended Data Fig. 7A). Confirmation of annotations was provided by spatial transcriptomics, which localised LZ and DZp/DZd transcriptomes to adjacent microanatomical regions within GCs in the spleen at 30 dpi^39^ (Extended Data Fig. 7B). By 21 dpi, the relative frequency of GC B cell subsets had stabilized (Fig. 4D), consistent with an equilibrium having been reached within GCs at a population level.

Of note, anti-malarial intervention had little qualitative effect on GC B cell composition, particularly between 21-42 dpi (Fig. 4D), despite the quantitative reductions in proportions of B cells displaying any GC B cell phenotype, a phenomenon validated by flow cytometric assessment with established markers^46,47^ (Extended Data Fig. 7C & D). These data suggested functional stability of GCs over time and after antimalarial treatment, which was supported by stable *Aicda* gene expression in DZ GC B cells (Fig. 4E). To formally test for GC function, we quantified SHM rates by comparing BCR sequences to germline. Firstly, as expected, we detected few mutations in BCRs of either naive follicular B cells or subsets not expected to engage in SHM (Fig. 4F, Extended Data Fig. 8). In contrast, GC B cells exhibited a progressive increase in median number of mutations summed across heavy and light chains, at a constant rate of 3.71 mutations per week, which remained unaltered (3.79 mutations per week) after anti-malarial treatment (Fig. 4F). Of note, SHM was detected as early as 10 dpi, a timepoint dominated by unmutated extrafollicular plasmablasts (Fig. 4B). Thus, mutational analysis supported GC functional stability over time and after antimalarial treatment by revealing constant SHM rates. Furthermore, SHM rates observed here were similar to those in previous studies of Ighv regions in murine lymph node GC B cells after influenza infection^25,48^ (Fig. 4G), further suggesting that SHM in our model of experimental malaria was not impaired relative to that of viral infection.

Having confirmed GC functional stability, we also noted that by 42 dpi GC B cells harboured BCRs with a wide spectrum of mutations, from 0-40 mutations per cell, with little skewing towards only highly mutated BCRs, as quantified by a progressive increase in the interquartile range of mutation counts over time (Fig 4H). Furthermore, although IgG1, IgG2b, IgG2c, IgG3 or IgA isotypes were largely retained over time and after treatment (Fig. 4F), a proportion of GC B cells was evident which expressed IgM and harboured 0-10 mutations (Fig. 4F and Extended Data Fig. 9). Taken together, our analysis revealed that once GCs are seeded, an immediate and progressive diversification of participating B cells ensues, mediated not only by a constant SHM rate, but also by the emergence and retention of unswitched cells with low numbers of mutations.

### Anti-malarial treatment exerts quantitative limits on GC output

Although GC B cell subset composition and SHM rates were unaffected by antimalarial treatment, it nonetheless elicited a quantitative drop in GC B cell frequency (Extended Data Fig. 5B-C). Hence, we next hypothesized that output from GCs was impaired by anti-malarial treatment. Consistent with this, inferred switched memory and atypical memory B cells, with mutations in BCRs consistent with GC origin (Extended Data Fig. 10), exhibited reduced numbers in the spleen following anti-malarial drug treatment, particularly at later stages of the infection (Extended Data Fig. 5C).

Secondly, we assessed GC output by searching for emerging plasma cells in the spleen, by re-clustering ASC transcriptomes within scRNAseq data of IgD^lo^ CD19^+^ B220^int-hi^ cells (Fig. 5A). This revealed a cluster expressing high levels of *Prg2,* a putative marker of splenic plasma cells^49^ (Fig. 5A). Given our sorting strategy had precluded detection of mature B220^-^CD19^-^ mature plasma cells, we reasoned splenic *Prg2*^hi^ ASCs, which had isotype-switched (Fig. 5B), and harboured high mutation counts from 21 dpi (Fig. 5C), were likely pre-plasma cells. *Prg2*^hi^ pre-plasma cell transcriptomes appeared to drop in frequency after anti-malarial treatment (Fig. 5D), providing further evidence of quantitative reduction in GC output. Finally, to formally screen for GC output we quantified long-lived plasma cells (LLPCs) amongst ASCs in the bone marrow, via flow cytometric assessment of CD138, TACI, CD19 and B220^50,51^ (Fig. 5E). At 42 dpi we detected a reduction in LLPCs in bone marrow after antimalarial drug treatment (Fig. 5E), suggesting GC output had been impaired. Consistent with reduced LLPC numbers, anti-malarial drug treatment also reduced circulating parasite-specific IgG (Fig. 5F) and rendered mice more susceptible to high-dose homologous re-challenge (Fig. 5G). Taken together, our data reveal that anti-malarial treatment exerts quantitative limits on GC output during experimental malaria.

**Figure 5:**
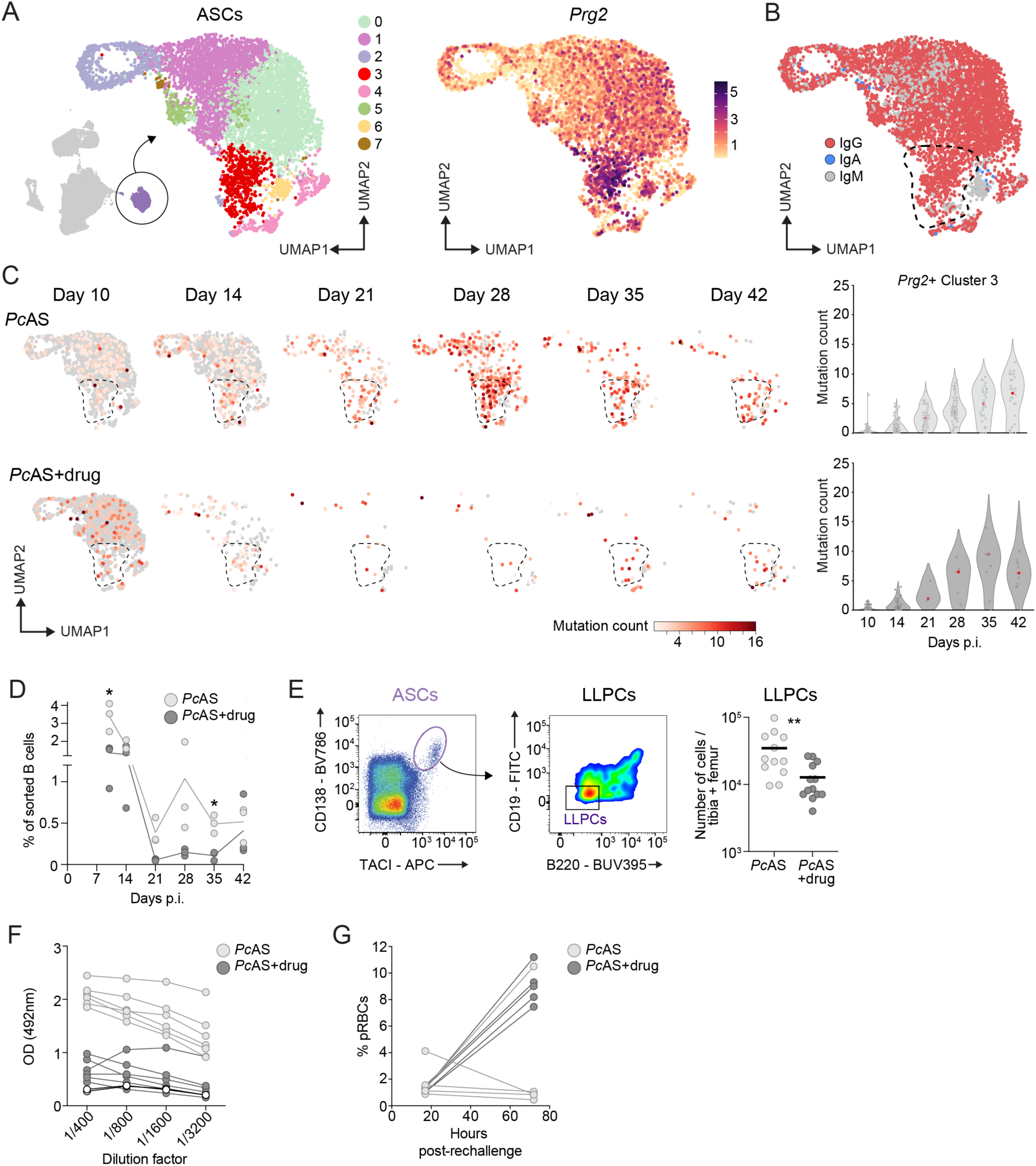
Anti-malarial treatment exerts quantitative limits on GC output. (**A**) UMAP of sub-clustered antibody secreting cells (ASCs) identifies *Prg2*^hi^ splenic plasma cells (red). (**B**) UMAP of isotype classification with *Prg2*^hi^ cluster indicated. (**C**) UMAPs of BCR mutation counts in ASCs per timepoint, with or without anti-malarial drug treatment. *Prg2*^hi^ area indicated by enclosed dashed line and quantified in violin plots (right), where the median number of mutations from violin plots are highlighted in red (**D**) Line graph of *Prg2*^hi^ cell frequencies (cluster 3 plasma cells) over the course of infection with or without anti-malarial drug treatment. (**E**) Representative pseudocolour density plots of ASCs and long-lived plasma cells (LLPCs) from day 42 *Pc*AS-infected bone marrow with scatter plot (right) of total LLPCs per tibia and femur. (**F**) Line graph of *Plasmodium*-specific IgG levels in the sera of mice at 60 dpi with or without anti-malarials as measured via ELISA (uninfected mice, white dots). (**G**) Mice treated with or without anti-malarials during a primary *Pc*AS infection were rechallenged and parasitemia in the blood measured 17 and 72 hours post-rechallenge. Lines in panel D connects the mean values at each timepoint and compared using Welch’s *t*-test. Data in panel E analysed by Mann-Whitney *t*-tests of *n* = 12-13 per group. Data are pooled from two independent experiments. Data in panel E represented by the mean. \**P* < 0.05, \*\**P* < 0.01.

### B cell lymphopoiesis relocates to the spleen during experimental malaria

Presence within GCs between 21-42 dpi of unswitched B cells harbouring few mutations suggested these as recent invader clones, consistent with viral infection and immunisation models^25,26^. In considering ongoing sources of naive B cells, we noted from 14 dpi, pronounced emergence of cells lacking *Ighd*, yet expressing *Il7r* and surrogate light chain gene *Vpreb1,* suggestive of precursor B cells (Extended Data Fig. 1C and 6A). To test for developing B cells in the spleen, we integrated our scRNAseq data (Extended Data Fig. 11A-C), with a dataset of developing B cells in the bone marrow^52^ (Extended Data Fig. 11D-E). Splenic data merged with bone marrow transcriptomes in certain expected areas, *e.g.* mature B cells (cluster 1) and ASCs (cluster 13) (Extended Data Fig. 11D), providing confidence in the integration. Inferred splenic immature and precursor B cell populations integrated with reported bone marrow counterparts (Extended Data Fig. 11D). Based on the integrated embedding and canonical markers, we annotated splenic precursor B cells as pro-and cycling pro-B cell (*Vpreb1, Igll1, Dntt, Rag1, Il7r, Il2ra, Ebf1, Ighd^-^, Cd74^-^, Ms4a1^-^*), pre-B cells (*Rag1, Il7r, Il2ra, Ebf1, Ighd^-^, Cd74^-^, Ms4a1^-^)* and immature B cells (*Ighd^-^, Ighm^+^, Ms4a1, Cd74*) on our integrated Experiment 1 and 2 UMAP (Fig. 6A and Extended Data Fig. 11F). Mapping transcriptomes over time suggested pro- and cycling pro- B cells emerged first at 14 dpi before subsiding, followed by pre-B cells peaking at 21 dpi, with immature B cells persisting until 42 dpi (Fig. 6A and Extended Data Fig. 12A). Anti-malarial drug treatment had little effect on these dynamics (Extended Data Fig. 12A). We then confirmed these dynamics via flow cytometric analysis of IL7R, IgM, CD43 and CD24 on IgD^lo^ CD93^+^ non-plasmablast, non-GC B cells (Fig. 6B and Extended Data Fig. 12B), and secondly that the phenomenon was unaffected by anti-malarial drug treatment (Extended Data Fig. 12C).

**Figure 6:**
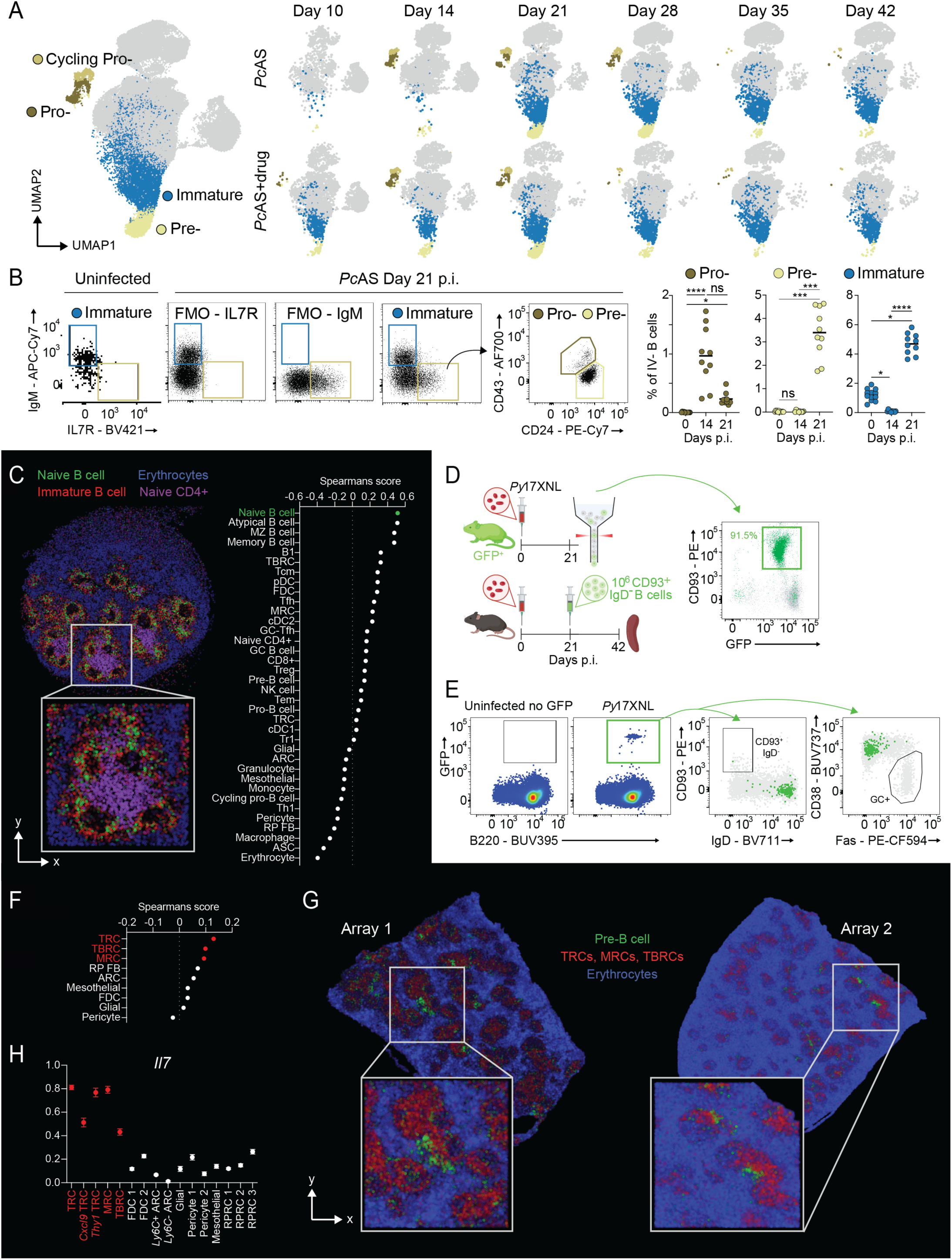
B cell lymphopoiesis relocates to the spleen during experimental malaria. (**A**) Annotated B cell scVI-integrated (Experiment 1 & 2) UMAP highlighting specific precursor B cell subsets (pro-, cycling pro-, pre- and immature B cells) split over the course of infection, with or without anti-malarial drug treatment. (**B**) IL7R and IgM surface markers were used to identify precursor B cells shown by representative dot plots of B cells gated on CD45.2^-^ (intravenous label negative, IV^-^), CD19^+^ B220^+^ CD138^-^ CD38^+^ Fas^-^ IgD^-^ CD93^+^ for immature B cells (IgM^+^ IL7R^-^), pro-B cells (IgM^-^ IL7R^+^, CD43^hi^ CD24^int^) and pre-B cells (IgM^-^ IL7R^+^, CD43^int^ CD24^hi^) in uninfected mice and at day 21 post-*Pc*AS infection, with quantitation of splenic precursor B cells at 0, 14 and 21 dpi. Gating determined by representative FMOs. (**C**) Robust Cell Type Deconvolution (RCTD) of immature B cell populations in a representative *Plasmodium* infected spleen at 30 dpi using *Slide-seqV*2, showing the spearman’s correlation score of immature B cell co-localisations with cell types from a scRNAseq reference across four independent 30 dpi spleen sections. (**D**) Sorting strategy for adoptive transfer of GFP^+^ splenic precursor B cells into *Py*17XNL infected host mice, showing enrichment of CD93^+^ GFP^+^ precursor B cells (right). (**E**) Representative pseudocolour density plots identifying GFP^+^ cells at day 21 post-transfer (green), compared to host-derived cells (grey). (**F**) RCTD spatial deconvolution of pre-B cell populations at 21 dpi using *Stereo-seqV1.2* spatial transcriptomics indicates spearman’s correlation scores of pre-B cell co-localisations with various splenic stromal cell types from a scRNAseq reference. Highlighted in red, T cell zone reticular cells (TRCs), T-B reticular cells (TBRCs) and marginal reticular cells (MRCs). (**G**) Location of pre-B cells, white-pulp stromal cells and erythrocytes on both replicate arrays following RCTD spatial deconvolution. (**H**) *Il7* gene expression across multiple splenic stromal cell types from a publicly available dataset^50^. Data in panel B represented by the mean and analysed by Kruskal-Wallis test with Dunn’s multiple comparisons test of *n* = 10-20 per group. Data in B are pooled from 2 independent experiments. \**P* < 0.05, \*\*\**P* < 0.001, \*\*\*\**P* < 0.0001. ns, not significant.

To investigate whether immature B cells might complete their development in the spleen, we first examined their location within the spleen at 30 dpi. Spatial transcriptomics at 10µm resolution, followed by cell-type deconvolution as previously^39^, revealed immature B cell transcriptomes co-localised with naïve B cells within follicles (Fig 6C, Extended Data Fig. 13A-B), consistent with potential to complete maturation in the spleen. To formally test this, we adoptively transferred splenic CD93^+^ IgD^-^ cells, also expressing eGFP, from infected mice (confirming splenic precursors at 21 dpi, Extended Data Fig. 14A) into infection-matched mice (Fig. 6D), and confirmed maturation 21 days later via loss of CD93 and acquisition of surface IgD (Fig. 6E). Of note, these cells had not given rise to ASCs or GC B cells by this time-point (Fig. 6E and Extended Data Fig. 14B). Together, these data demonstrated that splenic B cell precursors could traffic to follicles and complete maturation into naïve B cells amid an ongoing humoral immune response.

Finally, we sought to infer cellular and molecular mechanisms supporting B cell development in the spleen. High-resolution spatial transcriptomics at 21 dpi revealed non-random distributions of pre-B cells adjacent to white pulp zones, and, in particular, their co-localisation with multiple white pulp stromal cells: T cell zone reticular cells (TRCs), T-B reticular cells (TBRCs) and marginal reticular cells (MRCs) (Fig. 6F & G). Of note, these cell-types express high levels of *Il7* at steady state^53^ (Fig. 6H), a cytokine important for B cell development in the bone marrow^54-56^, with its receptor, *Il7r*, also expressed by splenic pre-B cells in our model (Extended Data Fig. 11F). Together, these data support a role for multiple white pulp stromal cell populations in supporting B cell development in the spleen, a process that temporally overlaps with SHM-mediated diversification of B cells in GCs.

### A Graphical User Interface (GUI) enables multi-parameter assessment of B cells *in vivo*

Having observed in our scRNAseq/BCRseq time-course a spectrum of overlapping B cell processes, including initial activation, CSR, clonal expansion, ASC differentiation, SHM in GCs, bystander activation, and splenic B cell lymphopoiesis, we next curated the data into a Graphical User Interface (GUI) alongside the high-resolution spatial transcriptomics data. (https://bcell-dynamics.science.unimelb.edu.au). The GUI enables simultaneous interrogation of individual B cells for 1) time-point, 2) cell-type 3) gene expression profile, 4) antibody isotype, 5) BCR sequence/mutation count, 6) clonal relationships with other cells, and 7) location of specific B cells states relative to other immune and non-immune cell types within the spleen. We reasoned such multi-parameter assessment would assist in hypothesis generation relating to B cells within secondary lymphoid organs. We present below three examples highlighting the utility of this GUI.

#### Case Study 1: Mining for pathogen-specific immunoglobulins

Monoclonal antibodies, developed against known *Plasmodium* surface antigens, have shown promise clinically for preventing malaria in humans (CIS43LS and L9LS Abs)^57-60^. BCRs harbouring specificity to unknown *Plasmodium* antigens might also be of potential use, *e.g.* for *de novo* immunoglobulin discovery pertaining to the field of malaria-prevention. Hence, we employed our GUI to search for BCRs with potential to bind *Plasmodium* antigens. We screened GC B cell transcriptomes (expressing *Aicda*) at later timepoints (28-42 dpi) for those switched to IgG isotypes, and with high numbers of mutations within their BCRs. Paired IGH and IGL sequences were downloaded, cloned, and recombinantly expressed (Fig. 7A). Subsequent ELISA-based analyses confirmed *Plasmodium-*binding, relative to similarly expressed negative control BCRs (Fig. 7A). Thus, our GUI likely harbours a collection of BCRs exhibiting specificity to as yet uncharacterised blood-stage *Plasmodium* antigens. Experimental and/or computational approaches may in the future determine precise antigen-specificities.

**Figure 7:**
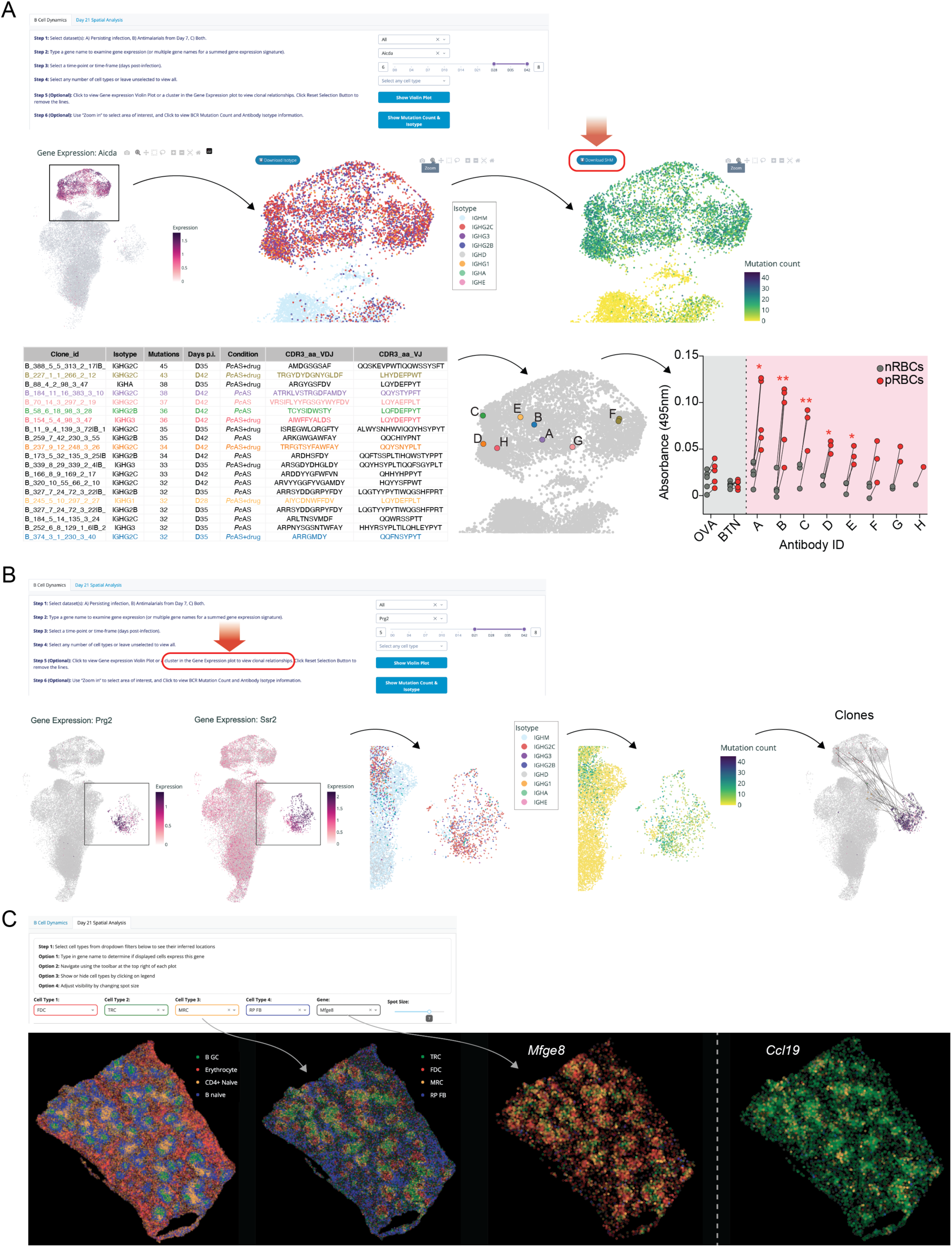
A Graphical User Interface (GUI) enables multi-parameter assessment of B cells *in vivo*. (**A**) Case study 1 – GC B cells screened for *Aicda* expression at 28-42 dpi and assessed for isotypes and mutation counts. Selected GC BCR data downloaded and sorted by mutation count. Coloured are BCRs that were cloned from the GC B cell dataset. ELISA measuring binding of cloned antibodies and irrelevant antibodies (OVA - ovalbumin and BTN - butyrophilin) to lysates from naive and parasitised RBCs. Data is shown as the mean ± S.D., *n* = 1-5 per antibody, dependent on antibody yield (A – 1.5mg, B – 1.5mg, C – 1mg, D – 0.8mg, E – 1mg, F – 0.09mg, G – 0.22mg, H – 0.18mg), analysed by multiple paired t-tests. \**P* <0.05, \*\**P* <0.01. (**B**) Case study 2 – ASCs screened for *Prg2^+^* and *Ssr2^-^* expression in 21-42 dpi and assessed for isotypes and mutation counts. Clicking on the ASC cluster highlights individual expanded clones. (**C**) Case study 3 – location of GC B cells, erythrocytes, naïve CD4^+^ T cells and naïve B cells assessed on *Stereo-seqV1.2* spatial transcriptomic datasets from day 21 *Pc*AS infected mouse spleens. Location of TRCs, FDCs, MRCs and red pulp fibroblasts (RP FBs), followed by assessment of these cell types expressing *Mfge8* or *Ccl19*.

#### Case Study 2: Inferring a pre-plasma cell state within the spleen

It was recently reported that differentiation towards a plasma cell fate in the spleen could be discerned via *Prg2* expression and concomitant down-regulation of *Ssr2*^49^. To further test this, we examined our GUI for evidence of ASCs expressing high levels of *Prg2,* and reduced expression of *Ssr2* (Fig. 7B). Having located these cells at 21-42 dpi, further evidence of these as pre-plasma cells was provided by having switched to IgG, harbouring high numbers of mutations consistent with having derived from the GC, and with clonal relationships with GC B cells (Fig. 7B). Hence, multi-parameter assessment at certain timepoints, of gene expression profiles, BCR isotype and mutation counts provided further of evidence of the existence of splenic pre-plasma cells.

#### Case study 3: Assessing cellular co-localisation and in situ gene expression within the spleen

To enable spatially-informed hypothesis generation regarding cell-type location, co-localisation and gene expression, we present within the GUI a genome-wide spatial transcriptomic (ST) data of two representative spleens at 21 dpi (when GCs are maturing, and B cell development is evident), generated from 1cm^2^ *Stereo-seqV1.2* arrays (with each pixel binned from 400 capture spots, approximating to a final working resolution of 10µm x 10µm) (Fig. 7C). This enables determination of relative location of several cell types (*e.g.* GC B cells, red blood cells (RBC), naïve CD4^+^ T cells and naïve B cells (Fig. 7C), as well as stromal cells. Displayed cell-types may then be filtered according to whether a given gene is detected at that location within the ST data. For example, having displayed four different stromal cell populations, TRC, FDC, MRC and RP FB, we observed *Mfge8* was expressed not only on FDCs as reported previously^61^, but also on other stromal cells such as TRCs (that also expressed *Ccl19*) and MRCs (Fig. 7C), which further support the notion that *Mfge8* expression is not limited to FDCs^62,63^.

Taken together, our GUI enables multiple parameter analysis of B cells in the spleen over time and space, via the examination of time points, cell-types, gene expression profiles, antibody isotypes, BCR mutation counts, clonal relationships, and spatial co-localisation.

## Discussion

Here we tested the hypothesis that population-level diversity amongst polyclonal B cell responses during infection is contributed to by intra-clonal diversification. We employed experimental malaria as a model, rather than single-antigen vaccination, because *Plasmodium* parasites express hundreds of proteins^29,30^, and might be expected to activate a broad spectrum of antigen-specific B cell clones. Furthermore, effective naturally acquired immunity to malaria benefits from a breadth of *Plasmodium*-specific antibodies^9,10^. By employing scRNAseq and paired BCRseq over time, we observed it was commonplace amongst hundreds of expanded clones, for the progeny of single naive B cells to vary both in the phenotypes and switched IgG sub-classes adopted. This suggests that during infection with a complex pathogen, although initial BCR sequence and antigen affinity can determine fate, as with single-antigen vaccination^22^, that BCR-independent processes may be important for diversifying the progeny of single naive B cells.

CSR can occur prior to GC formation during experimental vaccination^19-21^. Our observations during *Plasmodium* infection are consistent with this model, although the absolute timing of responses appears slower than that elicited by vaccination, perhaps due to low parasite numbers and hence antigen availability at early stages. scRNAseq over time facilitated the inference that *Myc-*upregulation, CSR and clonal expansion occur in a specific order, in which they nevertheless overlap with each other. Differential gene expression analysis presented here and visualised in our GUI highlights potential understudied genes in B cell activation, such as *Irf1*, which although implicated in gammaherpesvirus infection^64^, remains poorly studied in B cells. Recent work highlights methods for functional gene testing in naïve B cells^65^, which could be employed to test genetic targets mined from our data.

The consequences of temporal overlap between transcriptional programs associated with clonal expansion and CSR is that certain clones might commit to switching prior to clonal expansion, while others would initiate proliferation before committing to switching. Deeper sequencing of B cells from a single mouse raised the possibility that clonal isotype variegation may be the dominant outcome during infection, at least in our model, which further suggests substantial overlap and division-linked CSR *in vivo*, as suggested by previous *in vitro* studies^66-68^. To determine whether intra-clonal phenotypic variation is beneficial one might consider an alternative scenario in which initial BCR sequence fully determines fate and isotype in a given innate immune environment. In such a case, clones displaying specificity for an important, protective, pathogen-derived epitope would be locked into a single response that may not be optimal for pathogen control. In contrast, intra-clonal diversity early during infection may act as a “bet-hedging” strategy, providing responsive clones with functional flexibility, *e.g.* to enter the GC or not, to produce IgG2b, IgG3 or IgG2c, or to elicit rapid extra-follicular responses. Indeed, enhanced breadth of antibody isotypes mounted following malaria is associated with better clinical immunity^11^. Since many initial GC clones had also given rise to plasmablasts, we hypothesise GC entry remains an inclusive process that encourages diversity during recruitment for SHM.

Data from this study, together with previous reports^19-21^, prompts a consideration of why CSR initiates early, prior to GC entry. One explanation is that innate cytokine-mediated tailoring of humoral immune responses^69-71^, specifically of which antibody isotypes to deploy for a given pathogen class, must necessarily occur soon after pathogen sensing has occurred, *e.g* via pathogen-associated molecular patterns (PAMPs). This is likely to be within hours to a few days after pathogen exposure. Hence, it seems likely the CSR transcriptional program must be initiated at a time when innate immune responses have recently occurred, and innate cytokines are circulating within secondary lymphoid organs. Additionally, it may streamline affinity maturation within the GC by having many clones switched to IgG early during infection. Of note, this model does not preclude CSR from continuing within the GC, a prominent zone for AID expression^72,73^. Given the persistence of IgM^+^ GC clones in our model as in other studies^21,74^, we speculate that CSR continues in the GC across various times during infection.

The initial dynamics of SHM within GCs have been explored during hapten exposure or viral infection^25,48,75^. This has revealed linear rates of mutation acquisition in the heavy chains of BCRs of 1.9-2.1 mutations per week^25,48^. In our model, we also note a constant SHM rate over the first 6 weeks of infection of ∼2 mutations in the heavy chain per week, with similar rates evident for light chains. Our analysis suggests SHM initiates as early as 10 dpi during *Plasmodium* infection, at a rate comparable to viral infection^25,48^. Although our previous work, and from others revealed GC B cell responses are sub-optimal during *Plasmodium* infection^32,33^, we hypothesise this is due to limits on clonal expansion and fate choice, rather than adverse effects on SHM. We next consider why SHM rate is constant. AID expression levels are constant in GC B cells over time in our model. This is consistent with estimates that GC B cells mutate 1x10^-3^ per base pair per cell division^76,77^. However, although a recent study also suggested AID expression remained constant^78^, high affinity clones shielded themselves from deleterious mutations by spending less time during G0/G1 phase of the cell cycle, such that mutation rates were not constant amongst all GC clones^78,79^. Whether there is inter-clonal variation of SHM rates in our data is unknown. We also noted that anti-malarial treatment, which lowers parasite numbers and protects against splenomegaly, had no effect on SHM rates, despite deleterious effects on GC size, antibody titres, LLPC numbers and susceptibility during re-infection. Whether sufficient antigen remains on follicular dendritic cells (FDCs) to facilitate selection after anti-malarial treatment remains to be tested. Our data may be relevant for humans treated for mild malaria^31^, who could potentially harbour GCs in which affinity maturation continues unimpaired. However, complementary strategies with monoclonal antibodies or vaccines may be needed to quantitatively boost humoral immunity. In support of this, a recent phase III clinical trial suggests that combination of seasonal anti-malarial chemoprophylaxis and vaccination with RTS,S/AS01_E_ provide better clinical protection to malaria compared to either strategy in isolation^80^.

Our untargeted approach of examining polyclonal B cells facilitated the discovery of two phenomena that would have been missed with a focus on antigen-specific cells: Type I IFN-mediated bystander B cell activation, and a splenic B cell developmental pathway. Both these processes occurred contemporaneously, and in the same organ as antigen-specific B cell differentiation. Although long-term consequences of Type I IFN-mediated bystander activation of naïve B cells was not determined here, previous work including our own, indicates systemic blockade of Type I IFN-signaling early during an immune response improves cellular and humoral immunity to malaria^32,81-84^, and also during viral infection^85^. Whether Type I IFN-mediated effects on naïve B cells impairs subsequent immune responses to a new pathogen remains to be tested.

*Plasmodium* infection can impair haematopoiesis in the bone marrow, with negative consequences for myelopoiesis^86,87^. Common lymphoid progenitors are detected in adult mouse spleen^88^, with one study suggesting these could undergo B cell development during *Plasmodium* infection^89^. Our study extends this by revealing various immature B cell states are readily detected in the spleen during malaria, and that the classical B cell developmental pathway is likely phenocopied in the spleen. Further work is required to elucidate whether these mature B cells have the propensity to participate in ongoing or *de novo* immunological responses.

In summary, we have identified multiple, overlapping biological phenomena within splenic B cells during infection with blood-stage *Plasmodium* parasites. Amid a backdrop of bystander activation, single naive B cells with initial specificity for parasites proliferate and exhibit a spectrum of intra-clonal functional diversification. This diversity is maintained and increased in the GC via a constant rate of SHM, and persistence of both switched and unswitched GC B cells. Finally, deleterious effects on B cell development in bone marrow^90^ are counteracted by relocation to the spleen, potentially allowing the body to respond future immunological challenges.

### Limitations

Although our temporal assessment of polyclonal B cells aimed to capture the transition of B cells from one state to another, we were unable to identify an intermediate state between activated B cells and plasmablasts. This may have been due to the relative sparsity of timepoints at early stages of the immune response, rather than there being a lack of such an intermediate state. Secondly, the reduced capacity to detect BCR sequence data for GC B cells compared to plasmablasts prevented us from making unequivocal statements around the probabilities and frequencies with which naive B cells opted for these two fates. Rather, we restricted our conclusions to whether or not single clones giving rise to GC B cells could bifurcate into plasmablasts. Thirdly, our FACS-sorting strategy would not have isolated long lived plasma cells (LLPCs), meaning that plasma cell output from the GC could not be comprehensively assessed^91^. Instead, we infer *Prg2*-expressing ASCs in the spleen^49^ as likely pre-plasma cells yet to down-regulate CD19 as described by others^50^.

## Methods

### Mice

C57BL/6J mice were sourced from Animal Resources Centre (Perth, Western Australia, Australia) and maintained under specific pathogen-free (SPF) conditions in the Bioresources Facility (BRF) at The Peter Doherty Institute for Infection and Immunity (Melbourne, Victoria, Australia). Mice were housed at an ambient air temperature of 19-22°C and 40-70% humidity with a 12-hour day/night cycle. OB1.Rag−/−.uGFP, CD23Cre-MyD88flox and nzEGFP mice were bred in-house in the BRF of The Peter Doherty Institute for Infection and Immunity and maintained under the same conditions described above. Unless stated otherwise, all experimental mice used were female at 6-12 weeks old. All animal procedures were approved by the University of Melbourne Animal Ethics Committee (1915018 and 28191).

### Infection

Passage C57BL/6J mice were infected with a thawed vial of either *Plasmodium chabaudi chabaudi* clone AS (*Pc*AS) or *Plasmodium yoelii* strain 17XNL (*Py*17XNL) infected RBC stabilite. Once parasitemia reached 1-5%, blood was collected from passage mice by intra-cardiac bleed into RPMI containing 5U/ml heparin. A total of 10^5^ (*Pc*AS) or 10^4^ (*Py*17XNL) parasitized RBCs were injected into each experimental mouse via lateral tail vein injection. For rechallenge, mice were infected with 10^7^ parasitized RBCs via lateral tail vein injection, 28 days after primary infection with *Pc*AS.

### Parasitemia assessment

Parasitemia measurements were carried out as described previously^33^. Briefly, tail vein bleeds were performed to collect a drop of blood into RPMI containing 5U/ml of heparin. Diluted blood samples were stained at room temperature, for 30 minutes in the dark, with a combination of Hoechst 33342 (10µg/ml; Sigma-Aldrich) and Syto84 (5µM; Life Technologies). 10 times the initial volume of cold RPMI was used to quench the staining. The percentage of parasitized RBCs was determined as the proportion of Hoechst^+^ Syto84^+^ RBCs measured using a LSRFortessa X-20 cytometer (BD).

### Anti-malarial drug treatment

Artesunate (Guilin Pharmaceutical, kindly gifted by J. Moehrle) was made to 50mg/ml in 5% sodium bicarbonate solution (in PBS) and diluted with 0.9% saline (Baxter) to form a 5mg/ml solution of sodium artesunate. From day 7 post *Pc*AS infection, 1mg of sodium artesunate was injected intra-peritoneally (i.p.) twice daily to day 9, then once a day from day 10 to day 16 and twice weekly from day 17 to day 42 post *Pc*AS infection, alternating left and right side each i.p. injection. Additionally, from day 7 until day 42, mice received 70mg/l of pyrimethamine (Sigma Aldrich) in the drinking water.

### CD4+ T cell depletion

At 4 days prior to *Pc*AS infection, mice were administered 0.2mg of anti-CD4 depleting monoclonal antibody (clone GK1.5, BioXcell) or isotype control antibody (clone LTF-2, BioXcell) and a further 0.1mg dose was administered one day prior to infection. Antibodies were administered by intra-peritoneal injection in 200µl sterile PBS.

### IFNAR1 blockade

On the day of *Pc*AS injection, prior to infection, mice were administered 0.1mg of anti-IFNAR1 monoclonal antibody (clone MAR1-5A3, Leinco) or isotype control antibody (clone HKSP84, Leinco) and a further 0.1mg dose on day 2 post infection. Antibodies were administered by intra-peritoneal injection in 200µl sterile PBS.

### Intravenous Cell Labelling

To differentiate between lymphocytes that are in circulation and those that are within tissue, prior to spleen isolation, the mice were labelled with CD45.2 (clone 104, BioLegend) fluorescently conjugated antibody. A total of 15µl CD45.2-APC antibody was made up to 200µl in sterile 1x PBS and administered to each mouse via lateral tail vein injection. After 5 minutes, mice were culled and spleens harvested.

### Adoptive transfer

Uninfected spleens from OB1.Rag−/−.uGFP female donor mice were collected and homogenised through 70µm cell strainers. Homogenates were incubated with red blood cell (RBC) lysis buffer (Pharm Lyse, BD) for 3 minutes at room temperature. 15x10^6^ OB1 cells were injected into each C57BL/6J recipient mouse by lateral tail vein injection in a volume of 200µl.

For adoptive transfer of splenic precursor B cells, firstly, nzEGFP and C57BL/6J mice were infected with *Py*17XNL. At 21 dpi nzEGFP mouse spleens were collected, homogenised through 70µm cell strainers and underwent red blood cell lysis using Pharm Lyse (BD). Spleen homogenates were incubated with CD93-PE antibody (1:100) for 30 minutes on ice, washed twice with FACS wash buffer (2% FCS, 1mM EDTA in PBS) and then incubated with anti-PE MicroBeads (Miltenyi Biotec) (1:10) for a further 30 minutes on ice and then washed twice with FACS wash. Each homogenate was then passed through a LS MACS column as per the manufacturer’s instructions (Miltenyi Biotec, 130-042-401) for positive enrichment of CD93-PE cells. Enriched samples were pooled and cells were further sorted using a FACS Aria III (BD) to eliminate contaminating ASCs or GC B cells. Each infection-matched C57BL/6J recipient mouse received a total of 6-10x10^5^ GFP^+^ enriched splenic precursor B cells by lateral tail vein injection in a volume of 200µl. Mice were left for 21 days before assessment of adoptively transferred GFP^+^ cells.

### Flow cytometry

For bone marrow analysis, a single femur and tibia were isolated from each mouse. Bone marrow was extracted by flushing the inside of each bone with cold 1x PBS. Isolated bone marrow and spleens were homogenised through 70µm cell strainers and underwent red blood cell lysis using Pharm Lyse (BD). After washing in 1x PBS, cells were stained with Zombie Yellow live/dead stain (BioLegend) (1:500) in PBS on ice for 15 minutes, followed by Fc receptor block (1:500) (BD) in FACS wash buffer (2% FCS, 1mM EDTA in PBS) on ice for 20 minutes. Finally, cells were stained with different titrated panels of monoclonal antibodies diluted in FACS wash buffer (refer to table for antibody details). For intracellular staining, surface-stained cells were fixed and permeabilised using FoxP3/Transcription Factor Staining Set (eBiosciences) then stained with titrated antibody panels for 20-30 minutes on ice in the dark. Cells were acquired using LSRFortessa X-20 cytometer (BD), with all analysis of raw data performed using FlowJo version 5 (Treestar). Fluorescence minus one (FMO) controls for SCA-1, Ly6C, IgM, IL7R, CXCR4, CD69, CD83, CD86, CCR6, B220 and CD19 were used to help draw gates for each marker.

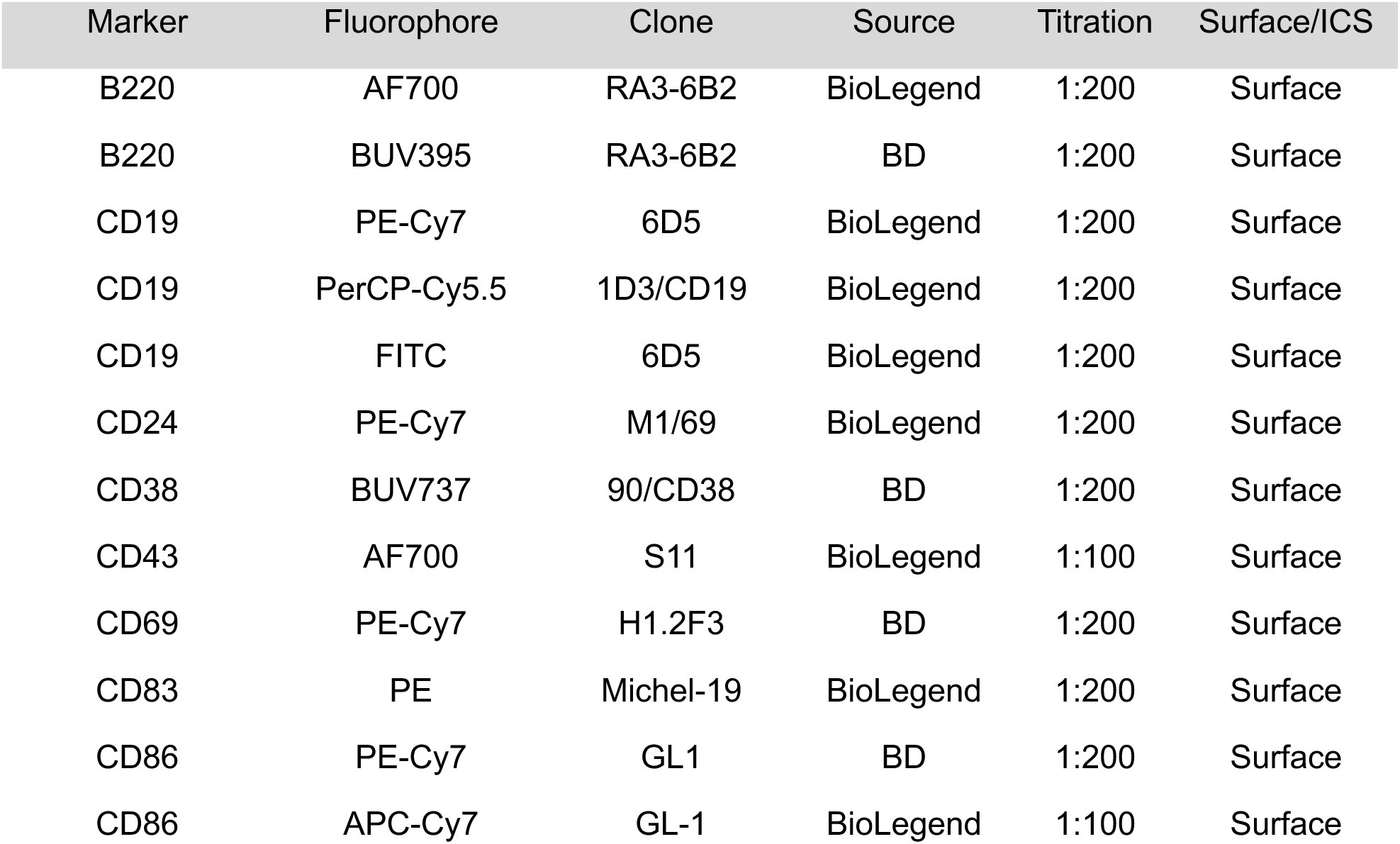
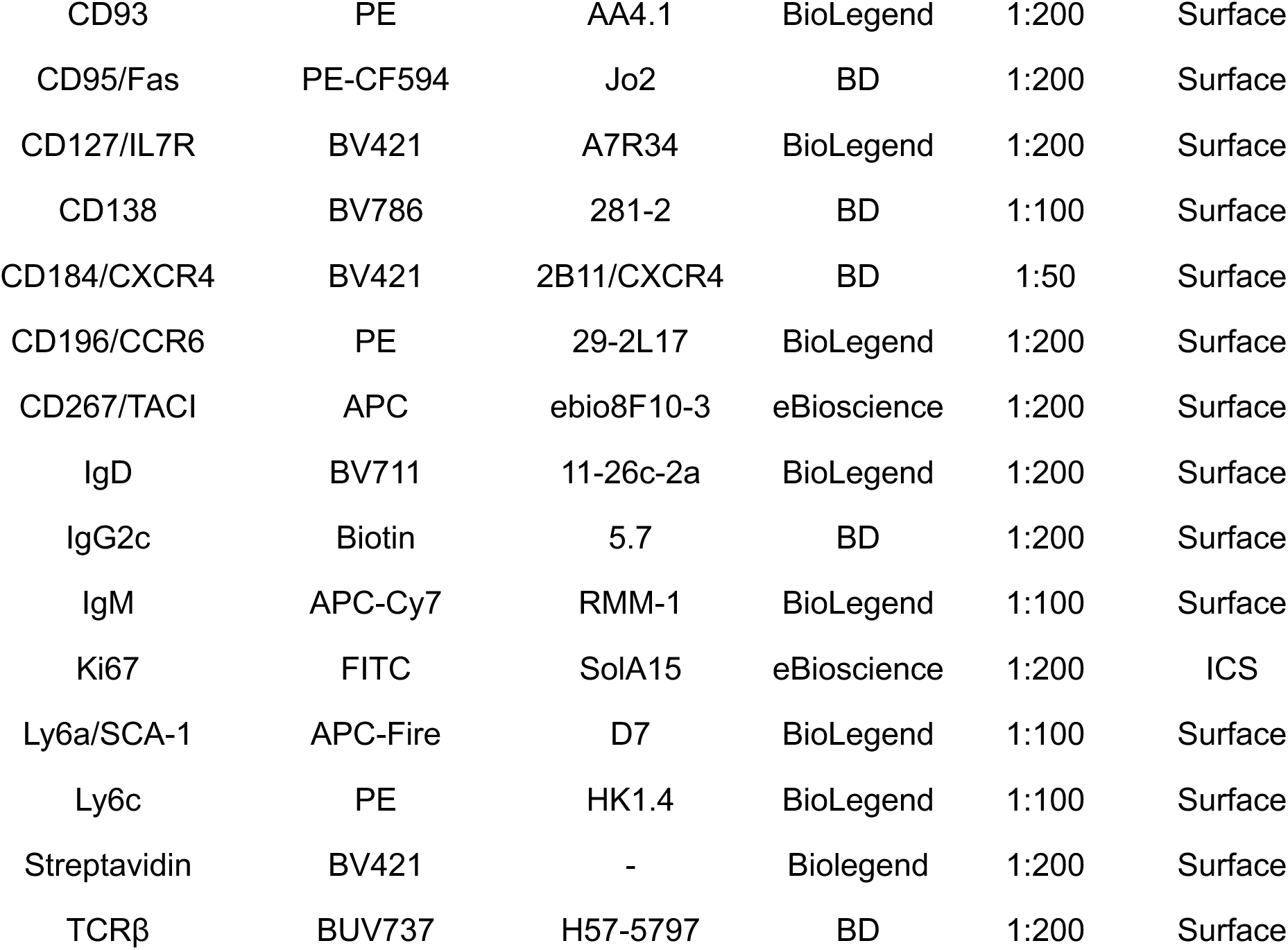

| Marker | Fluorophore | Clone | Source | Titration | Surface/ICS |
| --- | --- | --- | --- | --- | --- |
| B220 | AF700 | RA3-6B2 | BioLegend | 1:200 | Surface |
| B220 | BUV395 | RA3-6B2 | BD | 1:200 | Surface |
| CD19 | PE-Cy7 | 6D5 | BioLegend | 1:200 | Surface |
| CD19 | PerCP-Cy5.5 | 1D3/CD19 | BioLegend | 1:200 | Surface |
| CD19 | FITC | 6D5 | BioLegend | 1:200 | Surface |
| CD24 | PE-Cy7 | M1/69 | BioLegend | 1:200 | Surface |
| CD38 | BUV737 | 90/CD38 | BD | 1:200 | Surface |
| CD43 | AF700 | S11 | BioLegend | 1:100 | Surface |
| CD69 | PE-Cy7 | H1.2F3 | BD | 1:200 | Surface |
| CD83 | PE | Michel-19 | BioLegend | 1:200 | Surface |
| CD86 | PE-Cy7 | GL1 | BD | 1:200 | Surface |
| CD86 | APC-Cy7 | GL-1 | BioLegend | 1:100 | Surface |
| CD93 | PE | AA4.1 | BioLegend | 1:200 | Surface |
| CD95/Fas | PE-CF594 | Jo2 | BD | 1:200 | Surface |
| CD127/IL7R | BV421 | A7R34 | BioLegend | 1:200 | Surface |
| CD138 | BV786 | 281-2 | BD | 1:100 | Surface |
| CD184/CXCR4 | BV421 | 2B11/CXCR4 | BD | 1:50 | Surface |
| CD196/CCR6 | PE | 29-2L17 | BioLegend | 1:200 | Surface |
| CD267/TACI | APC | ebio8F10-3 | eBioscience | 1:200 | Surface |
| IgD | BV711 | 11-26c-2a | BioLegend | 1:200 | Surface |
| IgG2c | Biotin | 5.7 | BD | 1:200 | Surface |
| IgM | APC-Cy7 | RMM-1 | BioLegend | 1:100 | Surface |
| Ki67 | FITC | SolA15 | eBioscience | 1:200 | ICS |
| Ly6a/SCA-1 | APC-Fire | D7 | BioLegend | 1:100 | Surface |
| Ly6c | PE | HK1.4 | BioLegend | 1:100 | Surface |
| Streptavidin | BV421 | - | Biolegend | 1:200 | Surface |
| TCRβ | BUV737 | H57-5797 | BD | 1:200 | Surface |

### Antibody cloning

Selected BCR sequences (including ovalbumin, sequence adapted from^92^) were cloned into mouse IgG1 backbone expression vector for the heavy chain and mouse kappa backbone expression vector for light chain. Briefly:

1. Restriction Digestion: Cut heavy chain DNA and IgG1 vector with AgeI and SalI. Cut light chain mouse kappa light chain vector with AgeI and BsiWI (New England Biolabs). At 37°C for 2 hours, then run on an agarose gel.
2. Gel purification: DNA is extracted from the gel (Zymo Research D4001)
3. Ligation: Ligate each DNA fragment into corresponding expression vectors with T4 ligase (NEB M0202S) at 4°C overnight.
4. Transformation: Heat shock ligated DNA at 42°C for 90 seconds into DH5α competent cells
5. Miniprep plasmid purification (Zymo Research D4212)
6. Sequence verification by AGRF (Australian Genome Research Facility)
7. Maxiprep plasmid purification (Zymo Research D4202)

Antibodies were expressed by transient transfection into mammalian Expi293F cells using Expi-Fectamine (A14525, Thermo Fisher Scientific) with heavy and light chain plasmid (1:1 ratio) and purified by Protein G column chromatography (29048581 GE), followed by buffer-exchange into PBS. Butyrophilin antibody was synthesised previously^93^ as per clone sequence from^94^.

### Preparation of naive and parasitised RBC lysates

Parasitised antigen extract from infected blood was carried out as described previously^95,96^. Briefly, blood was collected from infected mice when blood parasitemia reached 20–30%. Blood was collected by intra-cardiac bleed into RPMI containing 5U/ml heparin. RBCs were washed once in RPMI at 400xg for 7 minutes at room temperature, before using ultrapure water to lyse, followed by four washes in ice-cold PBS at 16,000xg for 25 minutes at 4°C, followed by three cycles of freezing (two hours at -80°C) and thawing (30 minutes at room temperature). This procedure was followed by disruption (twice for 30s) with an ultrasonic probe at 60W on ice. Protein extracts were also processed from RBCs of uninfected C57BL/6J mice, to serve as negative controls in ELISA. Bradford assay (Thermo Fisher Scientific) was used to determine the concentration of proteins in the purified extracts.

### *Plasmodium-*specific IgG ELISA

As described previously^96^, Costar EIA/RIA 96-well flat bottom plates were coated overnight at 4°C with 10µl of parasite antigen (20µg) in bicarbonate coating buffer (pH9.6). Wells were washed three times with 0.05% Tween-20 in PBS and then blocked for 1 hour at 37°C with 1% skim milk in PBS. Wells were washed three times, followed by incubation with 5µg of purified cloned antibody for 1 hour at 37°C. Following six washes, wells were incubated in the dark with goat anti-mouse IgG HRP (G21040) for 30 minutes at room temperature. Wells were washed six times prior to development (100μl, OPD; P9187 Sigma-Aldrich) for five minutes in the dark before termination with an equal volume of 1M HCl. Absorbance was determined at 492nm using CLARIOstar analyser (BMG Labtech).

Serum was collected from uninfected and infected mice (day 55-60 post-*Pc*AS) with or without anti-malarial drug treatment and diluted 1/400, 1/800, 1/1600 or 1/3200 to assess binding to crude antigen extracts from *Pc*AS-infected pRBCs, as previously described^96^.

### Cell sorting and scRNAseq

Spleens were harvested from mice and homogenised through 70µm cell strainers. RBCs were lysed using Pharm Lyse (BD) for 3 minutes at room temperature and then single cell suspensions washed twice in 1x PBS. Samples were then incubated with Fc receptor-blocking antibodies (1:500) (BD) for 20 minutes on ice, followed by incubation with titrated monoclonal fluorescently conjugated antibodies and TotalSeq-C (BioLegend) mouse hashtags for 20 minutes on ice. Cells were washed twice and resuspended with propidium iodide (1:500) in PBS 2% BSA.

For experiment 1, samples from five individual mouse replicates were pooled together for each timepoint. Note, for timepoint day 4 post infection, there were four separately infected mice with one uninfected naive sample pooled in. Live, TCRβ negative, CD19^+^, B220^int-hi^ cells were sorted using a FACS Aria III (BD) into a 2% BSA/PBS buffer. For experiment 2, samples from seven individual mice (three for *Pc*AS, three for *Pc*AS+drug and one uninfected) were pooled together for each timepoint. Live, TCRβ negative, CD19^+^, B220^int-hi^, IgD^lo^ cells were sorted using a FACS Aria III (BD) into a PBS 2% BSA buffer. For each sample, a small number of IgD^hi^ naive cells were spiked in to assist in removal of batch effects for downstream scRNAseq analysis. For experiment 3, a single representative spleen was chosen from a 14 dpi mouse. Live, CD19^+^, B220^int-hi^ were sorted into either CD138^+^ IgD^-^ ASCs or CD38^-^ Fas^+^ GC B cells at 1:1 ratio. A small number of IgD^hi^ naive cells were spiked in.

After sorting, cells were loaded onto the Chromium controller and cDNA-sequencing libraries were prepared using the Chromium Single Cell 5’ Kits (10X Genomics).

### Analysis of scRNAseq datasets

For experiments 1 (days 0-14) and 2 (days 10-42), the transcriptome of splenic B cells was aligned using Cell Ranger 3.1.0 (10X Genomics) against mouse genome v2020-A (10X Genomics). For experiment 3, the transcriptome of splenic B cells was aligned using Cell Ranger 9.0.1 (10X Genomics) against mouse genome GRCm39-2024-A (10X Genomics). Hashtag demultiplexing and pre-processing of the data was performed using R version 4.2.0 (experiment 1 and 2) or version 4.3.1 (experiment 3) and Seurat version 4^97^. Doublets and negatives were removed from the demultiplexed samples for further downstream analysis (Supplementary Method Fig. 1A and 2A). For pre-processing, cells with a mitochondrial gene content of greater than 10% (experiment 1 and 2) or greater than 20% (experiment 3) were removed as well as cells with fewer than 200 and greater than 6000 total gene expression (Supplementary Method Fig. 1B and 2B).

Normalisation of data, selection of highly variable features and scaling of data were achieved using the *SCTransform()* function using default settings. Dimensional reduction was performed using the *RunPCA()* function and the resultant principal components were inputted to perform unifold manifold approximation and projection (UMAP) using *RunUMAP()* function. Finally, *FindNeighbours()* function and *FindClusters()* functions were used for unsupervised clustering.

Additionally, if immunoglobulin genes were left in the dataset for high dimensional reduction and clustering, clusters formed based purely on certain immunoglobulin variable heavy or light chain regions (Supplementary Method Fig. 1C and 2C). Therefore, immunoglobulin related genes (list of genes in Supplementary Table 2) were removed from the high dimensional reduction and clustering analysis, as recommended previously^98^. After high dimensional reduction and clustering analysis, UMAPs of each experiment were generated which highlighted anomalous cell clusters in each dataset: high mitochondrial content clusters, monocytes (*Lyz2*) and additionally in experiment 2, T cells (*Cd3g*) and erythrocytes (*Hbb-bs*) (Supplementary Method Fig. 1D and 2D). These clusters were removed from the UMAPs for subsequent downstream analysis. Altogether, after these quality control approaches, there were a total of 112,982 B cells: 40,623 B cells from experiment 1 and 72,359 B cells from experiment 2. A total of 19,482 post-QC B cell transcriptomes from a single mouse were retrieved from experiment 3.

### Cell type annotation

Differential gene expression analysis was performed using the *FindMarkers()* function with Wilcoxon rank sum test used to identify statistically significant differentially expressed genes across clusters. Geneset enrichment analysis (GSEA) was performed against biological processes GO terms using ToppGene Suite^99^ or WebGestalt^100^. Gene signatures were sourced from publicly available datasets for the following signatures: cell cycle^101^, class switch recombination (CSR)^27^, GC B cell subsets^46^ and B1a cells^102^. These cluster gene lists in addition to GSEA analysis and gene signatures aided in cell annotation of clusters for each UMAP.

### Sub-clustering

To further sub-cluster ASC and GC B cell subsets, each subset were taken separately and the above workflow repeated, i.e. normalisation, *SCTransform(), RunPCA(), RunUMAP(), FindNeighbours()* and *FindClusters()*.

### Data integration

Single-cell variational inference (scVI)^103^ was used to account for batch effects when performing data integration of day 10-42 B cell dataset with publicly available scRNAseq data of bone marrow B cells^52^. All parameters were kept at default and number of epochs set to 30 to train the model. Latent variables were used as input to run UMAP using Scanpy package (v1.9.1)^104^. In addition, data integration was performed on experiment 1 and 2 B cell datasets to generate a combined scRNAseq B cell dataset using the above parameters but using 9 epochs to the train the integration model.

### Bayesian Gaussian Process Latent Variable Model (bGPLVM)

Pseudo-temporal ordering of activated and GC B cells was performed using Bayesian Gaussian Process Latent Variable Model (BGPLVM) using GPFates^37^. This generated a progression of activated and GC B cell transcriptomes that correlated with sampled timepoints as well as an inferred pseudotime trajectory.

### Spatial transcriptomics using *Slide-seqV2*

Spatial transcriptomics was performed using *Slide-seqV2* at the Broad Institute of Harvard and MIT, as described elsewhere^105^. The methodology for the following spatial transcriptomic datasets were performed as previously described^39,106^ and analysis performed on previously published data^39^.

### Spatial transcriptomics using *Stereo-seqV1.2*

*Pc*AS infected spleens were isolated from 21 dpi mice and fresh frozen in liquid nitrogen, embedded in Tissue-Tek^®^ O.C.T Compound (Sakura) and stored at -80°C. *Stereo-seqV1.2* (BGI) was performed as per manufacturer’s instructions and two independent mouse spleens. Briefly, once appropriate red and white pulp regions were located within the spleen, a 10µm cryosection was mounted onto a spatially barcoded 1x1cm chip. A serial cryosection was collected to confirm spleen morphology via H&E staining. Spleen sections on chips were fixed with 4% PFA (BioScientific) followed by DAPI (Thermo Fisher Scientific) staining for 2 minutes. Chips were imaged using an Axioscan 7 microscope (Zeiss). Following imaging, tissue permeabilisation was conducted for 8 minutes as determined by preliminary permeabilisation tests. cDNA-sequencing libraries were prepared and sequenced data was processed following STOmics *Stereo-seq* Analysis Workflow (SAW) pipeline.

### Spatial deconvolution

Spatial deconvolution was achieved using robust cell type decomposition (RCTD) algorithm on day 21 *Pc*AS infected spleens (*Stereo-seqV1.2*) and on publicly available *Slide-seqv2* day 30 *Pc*AS infected spleens with artesunate treatment^107^ (n=4, two spatial transcriptomic arrays per spleen for two infected mice). The single cell transcriptomic references used were from the day 10-42 B cell dataset (experiment 2). In addition, our reference contained CD4, CD8 T cells, NK cells, macrophages, monocytes, erythrocytes, granulocytes, cDC1s, cDC2s and pDCs from an uninfected spleen reference dataset^106^ and GC-Tfhs from a day 30 infected *Pc*AS reference^39^. Finally, our reference contained cells from a publicly available spleen stromal dataset from uninfected mice, which included fibroblastic reticular cells (FRCs), follicular dendritic cells (FDCs), red pulp reticular cells (RPRCs) and adventitial reticular cells (ARCs)^53^. For cell-cell co-localisation analysis, we defined spatial neighbourhoods with a radius of 50µm. Spearman correlations are then calculated for all cell type pairs from the reference data and visualised as a heatmap.

### BCR sequencing and analysis

Dandelion package^42^ was used for VDJ clonal analysis.

### Pre-processing

Using Dandelion, the data was pre-processed using the standard outputs from all cellranger vdj versions. Here, single-cell V(D)J data from the 5′ Chromium 10X kit were initially processed with cellranger vdj pipeline (v6.1.2). BCR contigs contained in ‘all_contigs.fasta’ and ‘all_contig_annotations.csv’ files were then re-annotated using an immcantation-inspired pre-processing pipeline contained in the Dandelion singularity container (v0.3.0).

The pre-processing pipeline includes the following steps:

1. Adjust cell and contig barcodes by adding user-supplied suffixes and/or prefixes to ensure that there are no overlapping barcodes between samples.
2. Re-annotation of contigs with igblastn (v1.19.0) against IMGT (international ImMunoGeneTics) reference sequences.
3. Re-annotation of *D* and *J* genes separately using blastn with similar parameters as per igblastn (dust = ‘no’, word size ( J = 7; D = 9)) but with an additional *e*-value cutoff ( J = 10^−4^ in contrast to igblastn’s default cutoff of 10; D = 10^−3^). This is to enable the annotation of contigs without the *V* gene present.
4. Identification and recovery of nonoverlapping individual *J* gene segments (under associated ‘j_chain_multimapper’ columns). In the list of all mapped *J* genes (all_contig_j_blast.tsv) from blastn, the *J* gene with the highest score (j_support) was chosen. Dandelion then looks for the next *J* gene with the highest ‘j_support’ value, and with start (j_sequence_start) and end ( j_sequence_end) positions not overlapping with the selected *J* gene, and does so iteratively until the list of all mapped *J* genes are exhausted. In contigs without *V* gene annotations, it then selects the 5′ end leftmost *J* gene and update the ‘j_call’ column in the final AIRR table. For contigs with *V* gene annotations, but with multiple *J* gene calls, it uses the annotations provided by igblastn.

For BCRs, there are two additional steps:

1. Additional re-annotation of heavy-chain constant (C) region calls using blastn (v2.13.0) against curated sequences from CH1 regions of respective isotype class.
2. Heavy chain *V* gene allele correction using TIgGER (v1.0.0). The final outputs are then parsed into AIRR format with change-o scripts.

The rearrangement sequences that pass standard quality control checks are saved in file ending with the suffix ‘_contig_dandelion.tsv’.

### Postprocessing

In addition to the pre-processing steps at the contig level, postprocessing or integrating cell-level quality control is performed using Dandelion’s ‘check_contig’ function. The function checks whether a rearrangement is annotated with consistent *V*, *D*, *J* and *C* gene calls and performs special operations when a cell has multiple contigs. All contigs in a cell are sorted according to the unique molecular identifier (UMI) count in descending order, and productive contigs are ordered higher than non-productive contigs. For cells with other than one pair of productive contigs (one VDJ and one VJ), the function will assess if the cell is to be flagged with having orphan (no paired VDJ or VJ chain), extra pair(s) or ambiguous (biologically irreconcilable, for example, both BCRs in the same cell) status with an exception that IgM and IgD are allowed to coexist in the same B cell if no other isotypes are detected. The function also asserts a library type restriction with the rationale that the choice of the library type should mean that the primers used would most likely amplify only relevant sequences to a particular locus. Therefore, if there are any annotations to unexpected loci, these contigs likely represent artifacts and will be filtered away. A more stringent version of ‘check_contigs’ is used here by a separate function, ‘filter_contigs’, which only considers productive VDJ contigs, asserting a single cell should only have one VDJ and one VJ pair, or only an orphan VDJ chain, and explicitly removes contigs that fail these checks (with the same exceptions for IgM/IgD as per above). Productive contigs are selected for downstream analysis (Supplementary Methods Fig. 3A-B). Summary of frequency of unexpanded and expanded clones for each timepoint and experiment can be found in Supplementary Method Fig. 3.

### Clonotype definition and diversity

BCRs were grouped into clones/clonotypes using a sequential, rule-based procedure applied to productive rearrangements and implemented within Dandelion/Change-O outputs. Briefly, after cell-level QC (mentioned above), each cell contributed at most one dominant productive VDJ (heavy) and one productive VJ (light; IGK/IGL) contig (selected by UMI count), and clonotypes were assigned as follows:

1. **Chain-aware grouping (heavy and light):** Heavy- and light-chain repertoires were clonotyped separately using the criteria below, and paired clonotypes were defined using the combined heavy+light signature when both chains were available.
2. **Shared V/J gene usage:** Sequences were first required to have identical V-gene and J-gene calls (gene-level calls, not allele-level unless otherwise stated) for the relevant chain.
3. **Matched junction length:** Within each V/J bin, sequences were required to have identical CDR3 (junction) amino-acid length, ensuring a like-for-like comparison across aligned CDR3s.
4. **CDR3 similarity threshold:** Sequences passing (1–3) were then clustered by CDR3 amino-acid sequence similarity using a Hamming-distance criterion (equal-length CDR3s), with clonotype membership requiring ≥85% amino-acid identity.
5. **Handling incomplete receptors:** Cells lacking an unambiguous productive chain (e.g., missing light chain or orphan heavy chain) were clonotyped using the available productive chain only, and were excluded from analyses requiring paired heavy+light definition.

Diversity was quantified from the resulting clonotype frequency tables (counts of cells per clonotype within each sample/timepoint/condition). Clonal expansion was summarised by the distribution of clone sizes (e.g., unexpanded singletons versus expanded clones), and repertoire diversity was computed using standard clonotype-based diversity metrics (e.g., Shannon entropy and/or Simpson diversity) on depth-normalised clonotype tables to enable fair comparisons across samples.

### Somatic Hypermutation (SHM)

The basic mutation load is quantified by using the ‘pp.quantify_mutations’ in Dandelion which is a wrapper of SHaZam’s basic mutational analysis. It sums all mutation scores (heavy and light chains, silent and replacement mutations) for the same cell.

### GC B cell mutation analysis

Mutation rate analysis is calculated by fitting linear regression line to the median number of mutations at each timepoint for each condition (*Pc*AS or *Pc*AS+drug). The gradient of the line provides the mutation rate per day. Mutation rate analysis was also performed specifically on the Ighv region of the BCR in comparison with publicly available clonal data from lymph node GC B cells after influenza infection^25,48^.

### GUI Architecture

An interactive B cell GUI was developed using Dash (pyDash), which couples a Flask-based Python back end to a React-based front end, with all layouts and interactions defined in a single Python codebase. Interactive visualisation was implemented with Plotly, enabling hover-based inspection, zooming, brushing/linked filtering, and export across modules (B cell dynamics, gene expression, violin plots, clonality, and spatial views). Single-cell RNA-seq and paired BCRseq data were curated into an AnnData/HDF5 data model with harmonized metadata (treatment, timepoint, cell type, clonotype, isotype and SHM metrics).

To minimise runtime costs, 2D/3D UMAP embeddings and key summary metrics were pre-computed, and frequently queried metadata fields were indexed to support rapid subsetting in callbacks. For usability at scale, lightweight aggregation was applied where appropriate and the dataset was down-sampled explicitly retaining cells with BCR information to maintain a smooth, responsive user experience.

### Plotting

For data visualisation, the following packages were used: Prism version 9, Tableau (2023.2.0), Python version 3.8.8 or ggplot2 version 3.4.4 package utilised in R version 4.2.1. Adobe Illustrator version 26.3.1 was used for figure generation and schematics were partly generated using BioRender.

## Data availability

Raw sequencing data for scRNAseq B cell data for experiments 1, 2 and 3 and two independent *Stereo-seqV1.2* spleen samples are found via accession number provided: GSE286215 and can be accessed using reviewer’s token *cnaduagirzinval*. Processed data including contig annotations of BCR data, Seurat objects of all experiments and integrated Seurat object are also found via accession number: GSE286215 and can be accessed using reviewer’s token *cnaduagirzinval*. Previously published data used in this present study can be found here: spleen stromal scRNAseq data for spatial deconvolution (GSE171124), T cell scRNAseq data for spatial deconvolution (GSE244806, GSE233713), *Slide-seqV2* spatial transcriptomics data (GSE234253), B cell bone marrow scRNAseq data (GSE168158).

## Code availability

Publicly available software was used, as detailed in the methods and reporting summary. Code can be provided upon request.

## Ethics declarations

The authors declare no competing interests.

## Acknowledgements

This research was supported by The University of Melbourne’s Research Computing Services, the Petascale Initiative, and the LIEF HPC-GPGPU Facility hosted at The University of Melbourne. The facility was established with the assistance of LIEF grant LE170100200. We extend our gratitude to the Bioresources Facility located at The Peter Doherty Institute for Infection and Immunity for their animal care and assistance. We are also deeply thankful to the Doherty Institute node of the Melbourne Cytometry Platform at The University of Melbourne. We acknowledge the Hudson Genomics Facility for sequencing of single cell data. We would also like to acknowledge Dr Katherine Jackson (Garvan Institute of Medical Research, Sydney, Australia) for discussions and help regarding BCR sequencing analysis. A thank you also goes towards Dr Nick (Ka Leung) Li and Dr Jiyoti Verma from Decode Science for assistance and support with the *Stereo-seqV1.2* experiments as well as the Biological Optical Microscopy Platform at the University of Melbourne. The authors acknowledge the South Australian Genomics Centre (SAGC) for sequencing the *Stereo-seqV1.2* libraries. The SAGC is supported by the National Collaborative Research Infrastructure Strategy (NCRIS) via Bioplatforms Australia. We finally acknowledge Joshua Lilley at The University of Melbourne for assistance with deployment of the GUI that accompanies this resource.

## Extended Data Figures

**Extended Data Figure 1:**
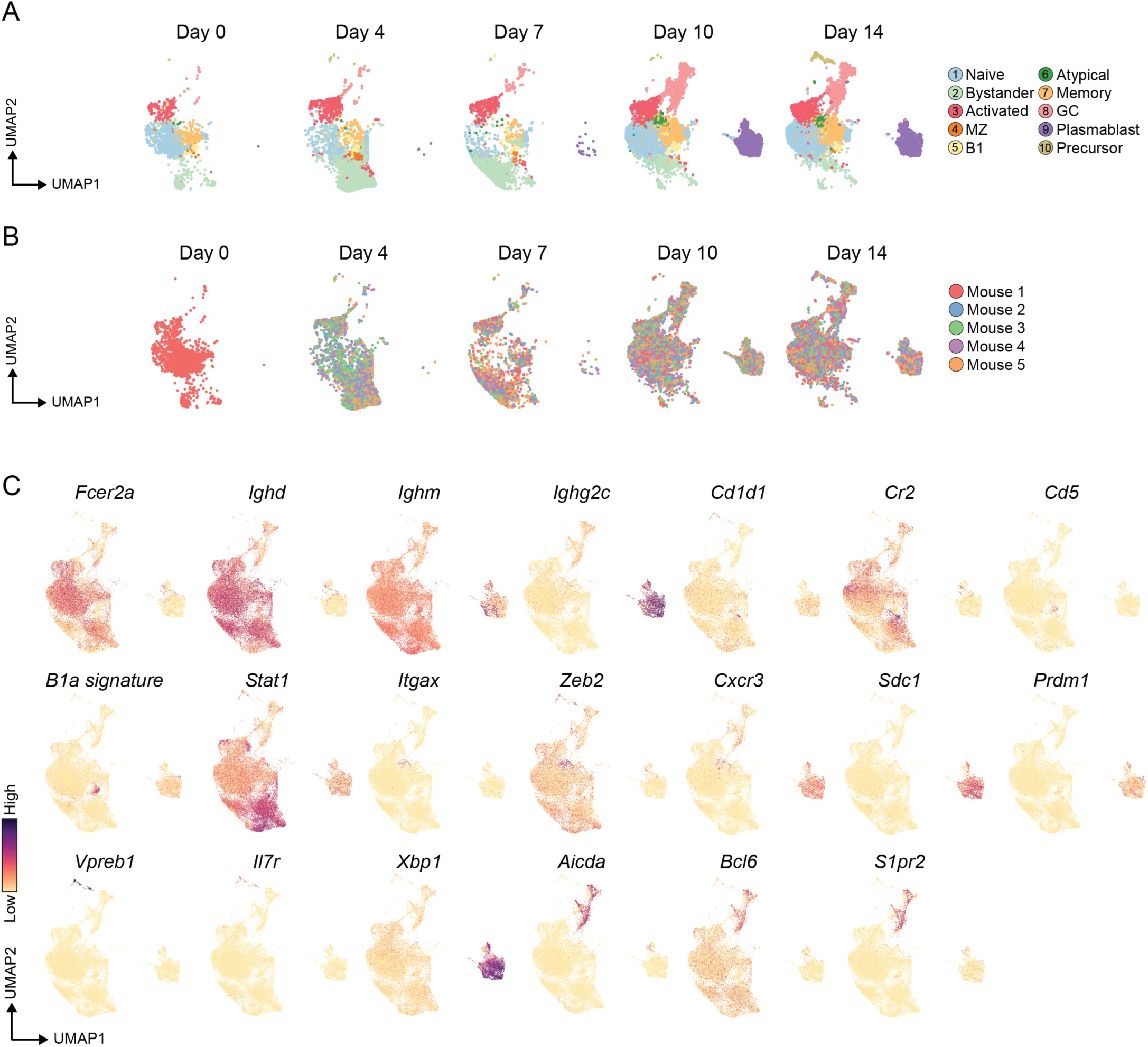
Cluster annotation of B cell scRNAseq data (Experiment 1: 0-14 dpi). (**A**) Annotation of B cell subsets after high-dimensional reduction and clustering analysis, split by timepoints and (**B**) grouped by individual mice. (**C**) Expression of canonical markers for each cluster type: naive follicular B cells (*Fcer2a*, unswitched: *Ighd^+^, Ighm^+^, Ighg2c^-^*), MZ cells (*Cd1d1, Cr2*), B1 cells (*Cd5, B1a signature*), bystanders (*Stat1*), precursor B cells (*Vpreb1, Il7r*), atypicals (*Itgax, Zeb2, Cxcr3*), plasmablasts (*Sdc1, Prdm1, Xbp1*, switched: *Ighd^-^, Ighg2c^+^*) and GC B cells (*Aicda, Bcl6, S1pr2,* switched: *Ighd^-^, Ighg2c^+^*).

**Extended Data Figure 2:**
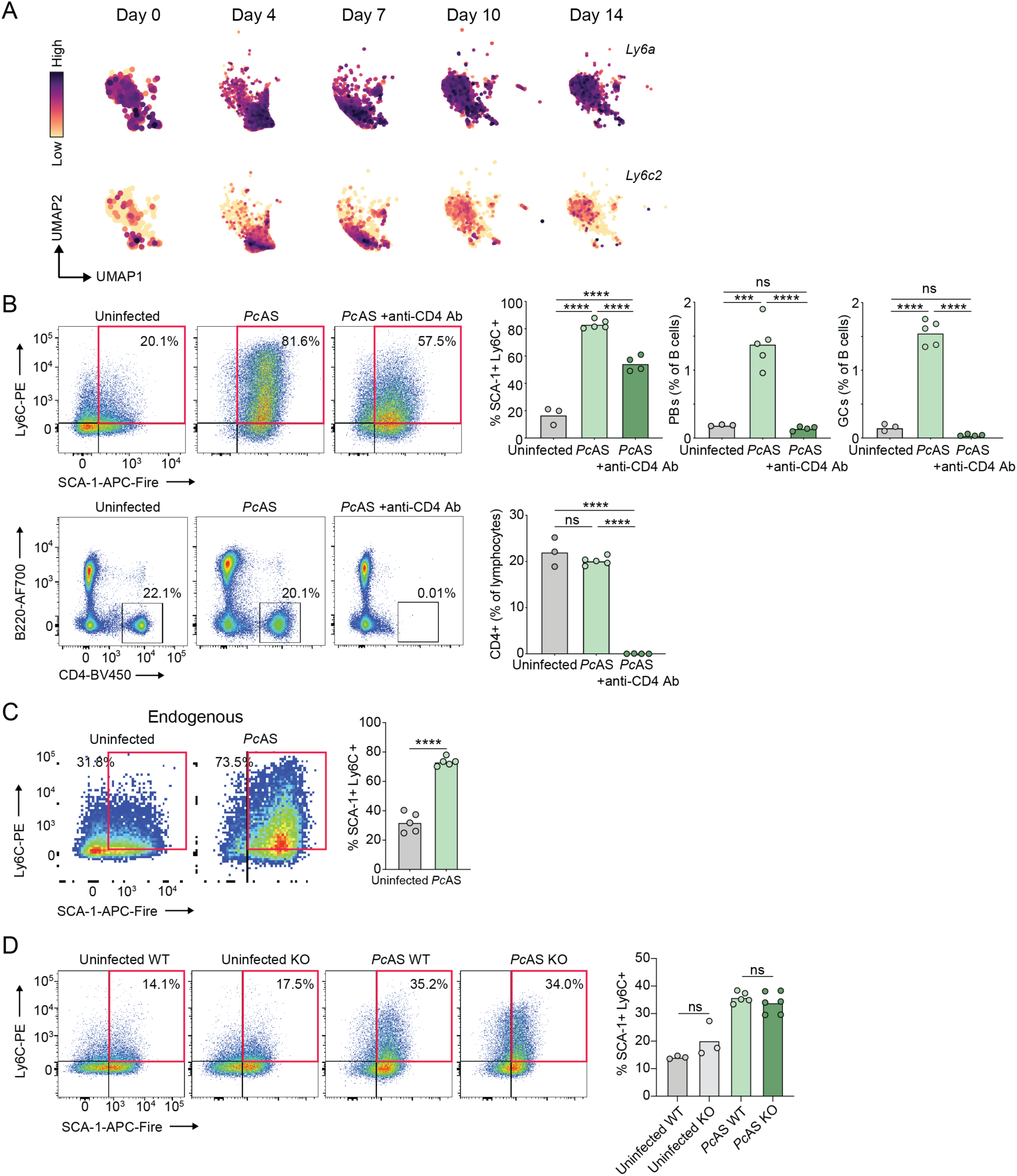
Bystander B cell activation is partially CD4^+^ T cell-dependent, and independent of cell-intrinsic MyD88-signalling. (**A**) Expression of *Ly6a* (Sca-1) and *Ly6c2* on naive follicular and bystander B cells over the course of *Plasmodium* infection. (**B**) Representative density dot plots and bar graphs of CD4 expression on lymphocytes 7 days post-*Pc*AS infection, and SCA-1 and Ly6C expression on IgD^+^ B cells, and frequency of plasmablast and GC B cells relative to all B cells 7 days post *Pc*AS infection, with or without CD4 T cell depletion. (**C**) Representative density dot plots and bar graph showing proportion of SCA-1 and Ly6C expression on CD86^lo^, IgD^+^ endogenous host derived B cells from the OB1 adoptive transfer experiments, 4 days post-*Pc*AS infection. (**D**) Representative density dot plots, and bar graph showing proportion of SCA-1 and Ly6C expression on IgD^+^ MyD88flox/flox B cells with (CD23^cre+/-^ KO) or without (CD23^cre-/-^ WT) MyD88 KO, 7 days post-*Pc*AS infection. Data in panels B-D are represented as the mean value. Data in panel B and D analysed using one-way ANOVA with Tukey’s multiple comparisons test and data in panel C analysed using Welch’s *t*-test. All data are representative of two independent experiments of (B & C) *n* = 3-5 and (D) *n* = 3-7 biological replicates per group. \*\*\**P <* 0.001, \*\*\*\**P* < 0.0001. ns, not significant.

**Extended Date Figure 3:**
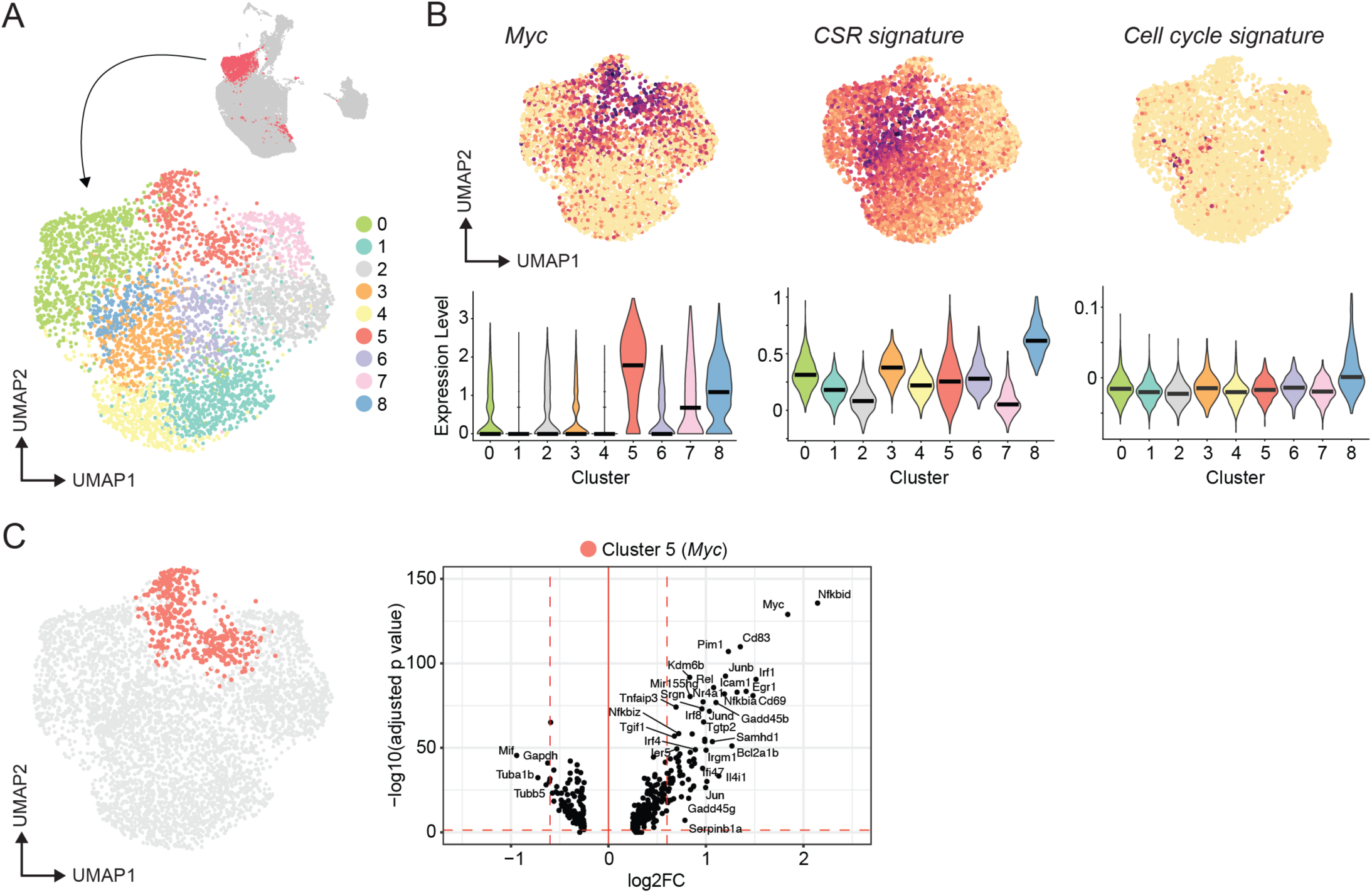
Re-clustering and deeper analysis of the early activated B cell cluster. (**A**) UMAP of sub-clustered activated B cells. (**B**) Expression of overlapping events of *Myc*, CSR and cell cycle signatures within each cluster. (**C**) Volcano plot of differentially regulated genes for Cluster 5 highlighting selected top differentially regulated and significant genes.

**Extended Data Figure 4:**
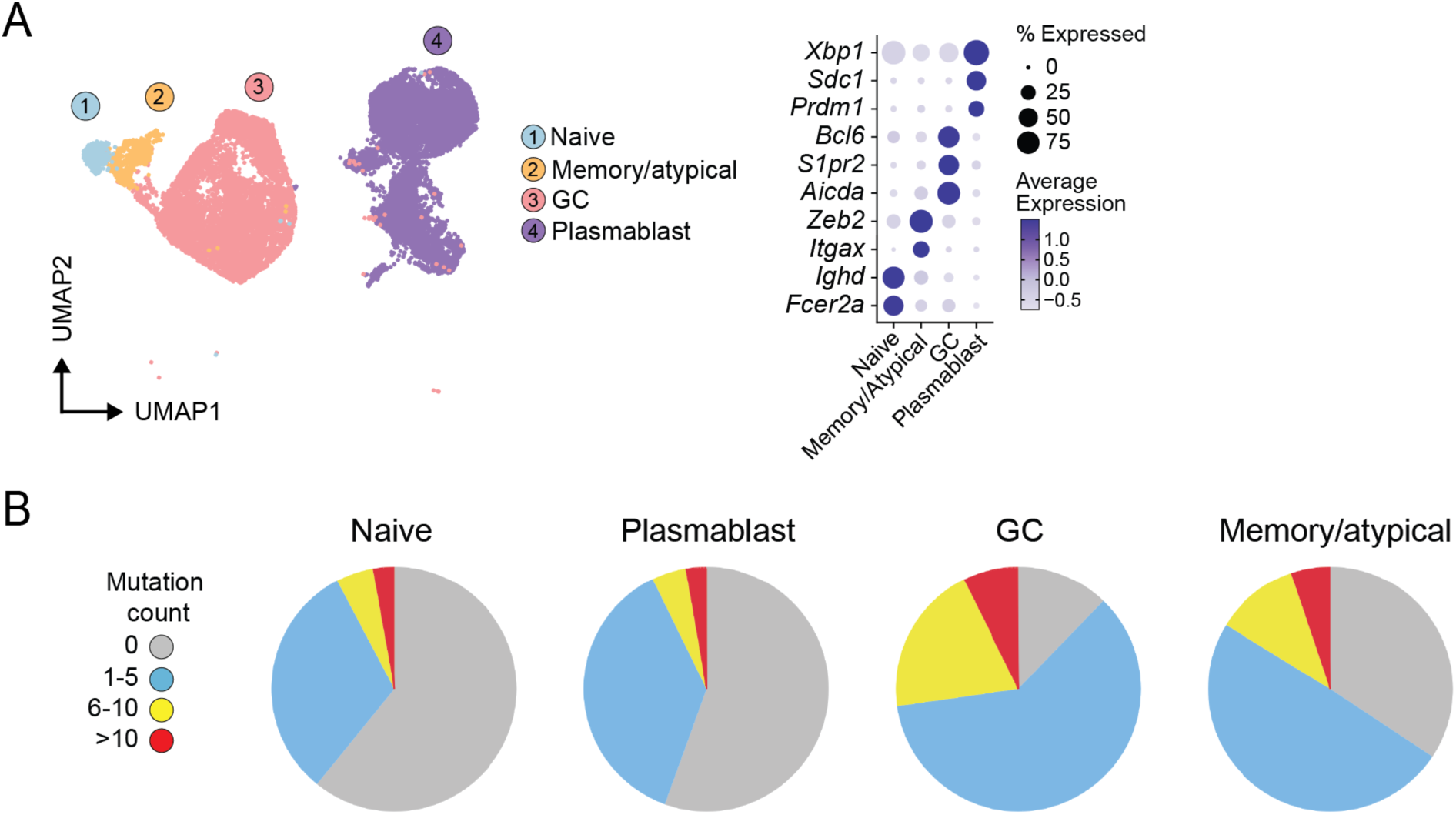
QC assessment of scRNAseq/BCRseq data from enriched GC B cells and plasmablasts from a single representative mouse spleen at 14 dpi. (**A**) Annotated UMAP of scRNAseq/BCRseq data generated from FACS-sorted splenic GC B cells, plasmablasts, minor populations of atypical/memory B cells, and naïve spike-in cells from a representative mouse at 14 dpi, with dot plot showing canonical gene markers. (**B**) Pie charts showing mutation count of clones detected in naïve B cells, GC B cells and memory/atypical B cells.

**Extended Data Figure 5:**
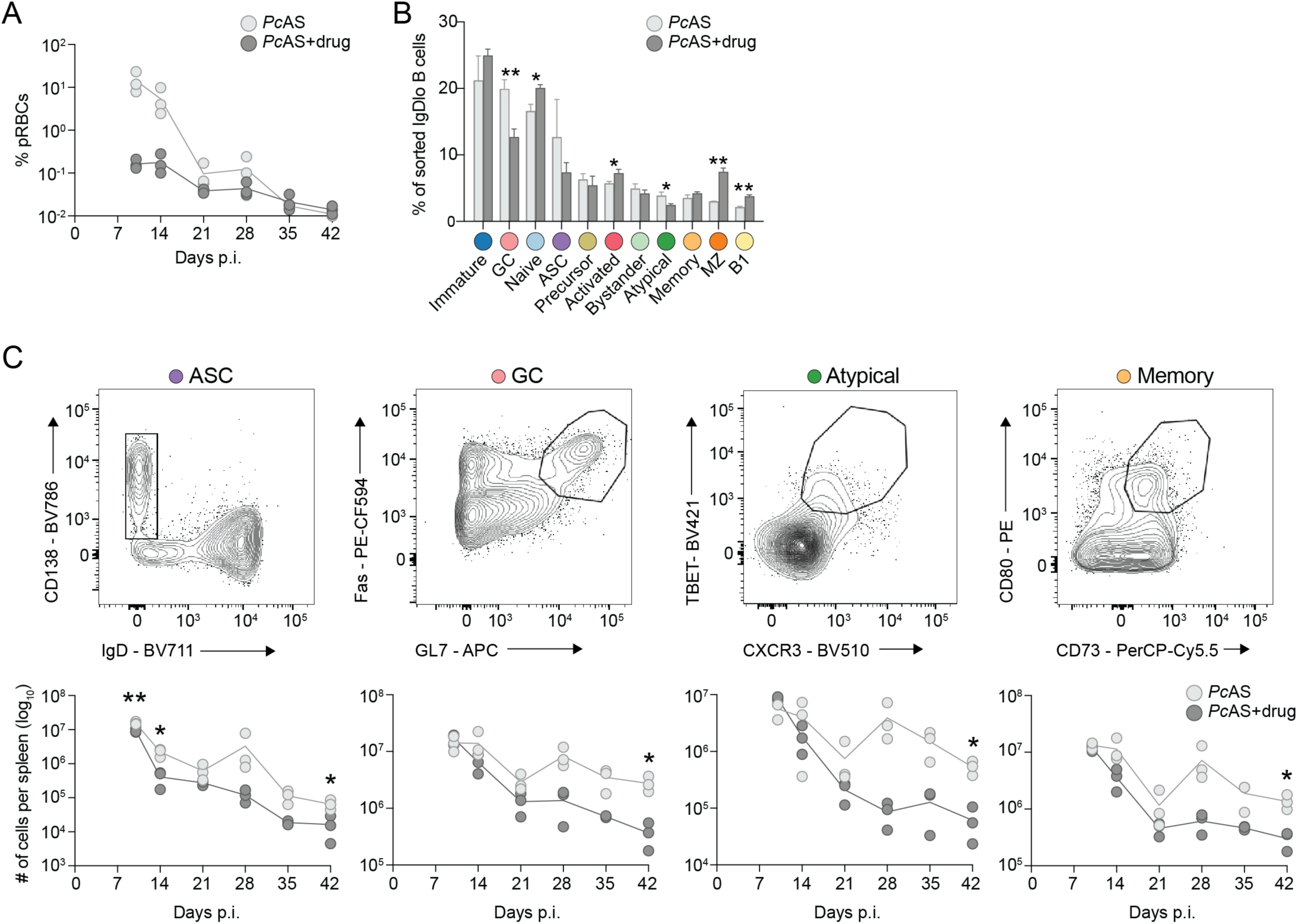
Anti-malarial treatment efficacy, and effects on proportions of B cell subsets in scRNAseq and FACS validation datasets. (**A**) Percentage parasitemia of circulating RBCs at each timepoint post with or without anti-malarials. Lines connect the mean values at each timepoint. (**B**) For the combined day 10-42 scRNAseq dataset, proportion of each B cell subset under each condition (*Pc*AS or *Pc*AS+drug) for individual mice. (**C**) Flow cytometric assessment of live CD19^+^ B220^int-hi^ cells for ASCs, GCs, atypical and memory B cells on matched spleen samples from the scRNAseq dataset at each timepoint post-infection, Data are the mean ± S.D, statistical test used was Welch’s *t*-test. \**P* <0.05, \*\**P* <0.01.

**Extended Data Figure 6:**
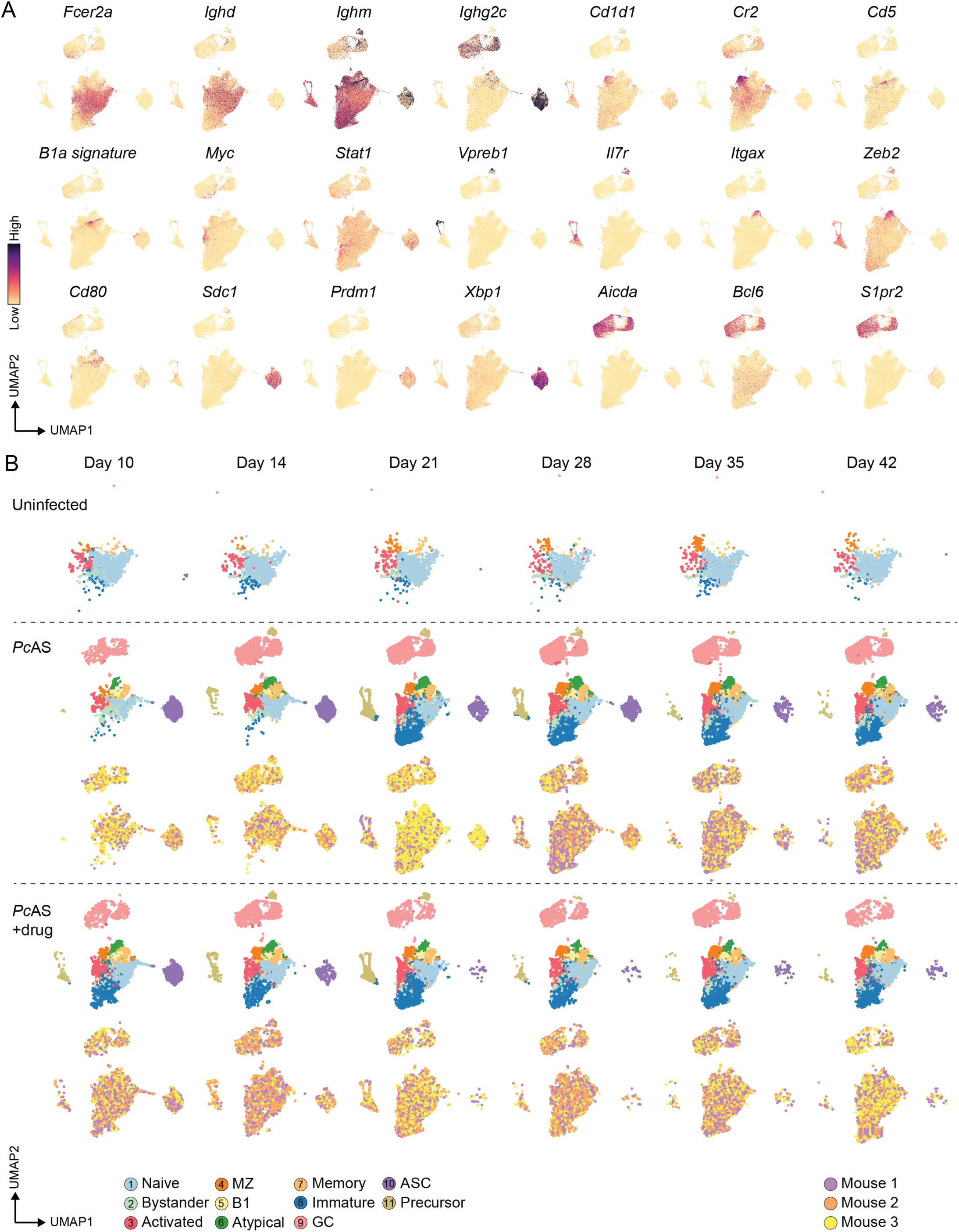
Cluster annotation of B cell scRNAseq data (Experiment 2: 10-42 dpi). (**A**) Expression of canonical markers for each cluster type: naive follicular B cells (*Fcer2a*, unswitched: *Ighd^+^, Ighm^+^, Ighg2c^-^*), MZ cells (*Cd1d1, Cr2*), B1 cells (*Cd5, B1a signature*), activated B cells (*Myc*), bystanders (*Stat1*), precursor B cells (*Vpreb1, Il7r*), atypicals (*Itgax, Zeb2*), memory B cells (*Cd80*), plasmablasts (*Sdc1, Prdm1, Xbp1*, switched: *Ighd^-^, Ighg2c^+^*) and GC B cells (*Aicda, Bcl6, S1pr2,* switched: *Ighd^-^, Ighg2c^+^*). (**B**) Annotated UMAPs split by timepoint for each condition: uninfected, *Pc*AS or *Pc*AS+drug also grouped by individual mice.

**Extended Data Figure 7:**
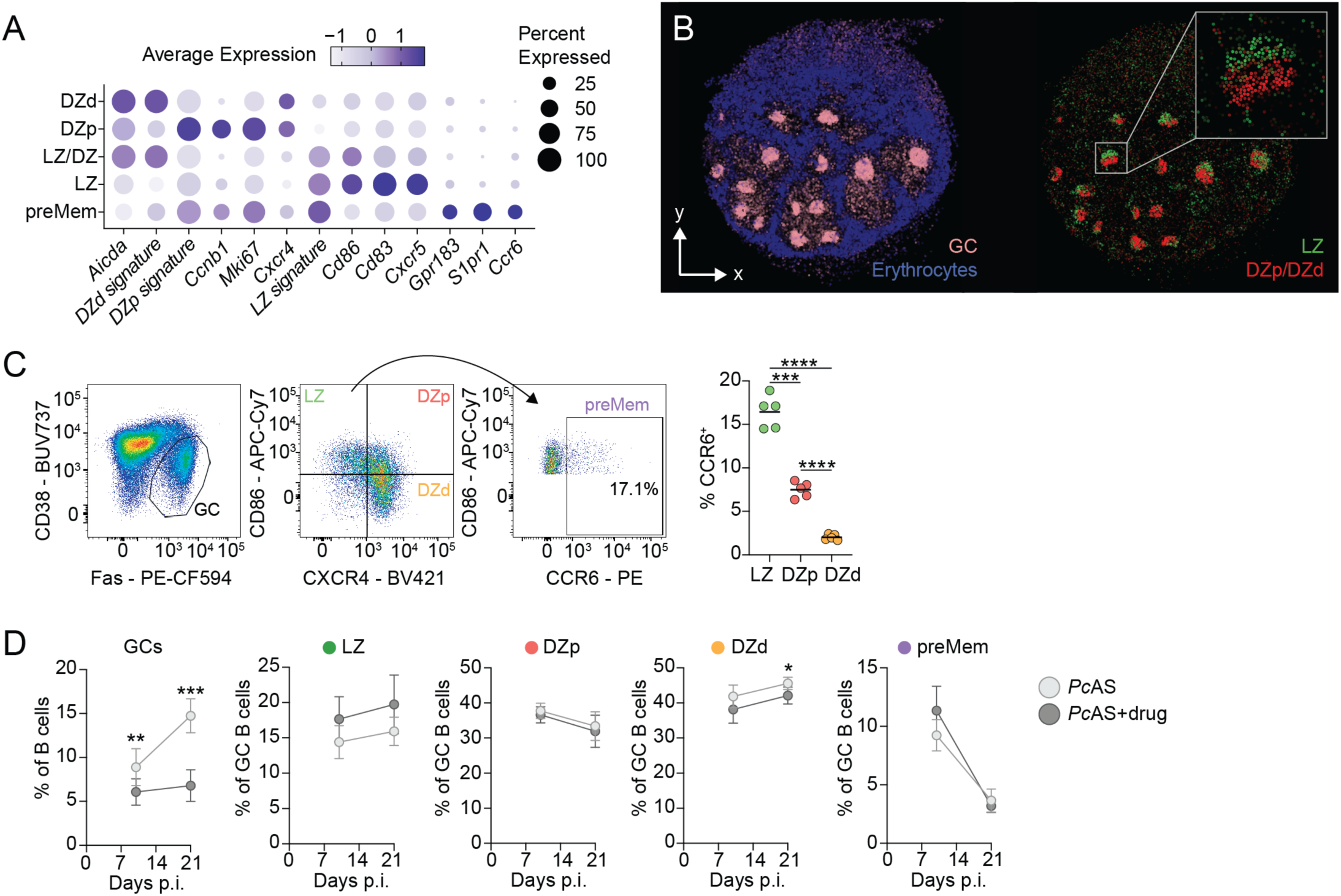
Anti-malaria drug intervention does not affect GC B cell composition. (**A**) Dot plot of canonical gene markers and signatures for LZ and DZ GC B cells and preMems. (**B**) RCTD spatial deconvolution of GC B cell populations on a day 30, *Plasmodium* infected spleen using *Slide-seqV2* spatial transcriptomics. (**C**) Gating strategy to identify CD38^-^ Fas^+^ LZ, DZp, DZd and preMem GC B cells in 21 dpi mouse spleens, showing quantitation of % CCR6+ in each GC B cell compartment. Quantitation shown is representative of two independent experiments. Data is mean, n=5 biological replicates, analysed using paired one-way ANOVA with Geisser-Greenhouse correction. (**D**) Quantitation of splenic GC B cells and LZ, DZp, DZd and preMem GC B cells in 10 and 21 dpi mice with or without drug treatment. Data is shown as mean ± S.D. pooled form two independent experiments, n=9-10 biological replicates, and analysed using multiple Welch’s t-tests. \**P* < 0.05, \*\**P* <0.01. \*\*\*\**P* <0.0001.

**Extended Data Figure 8:**
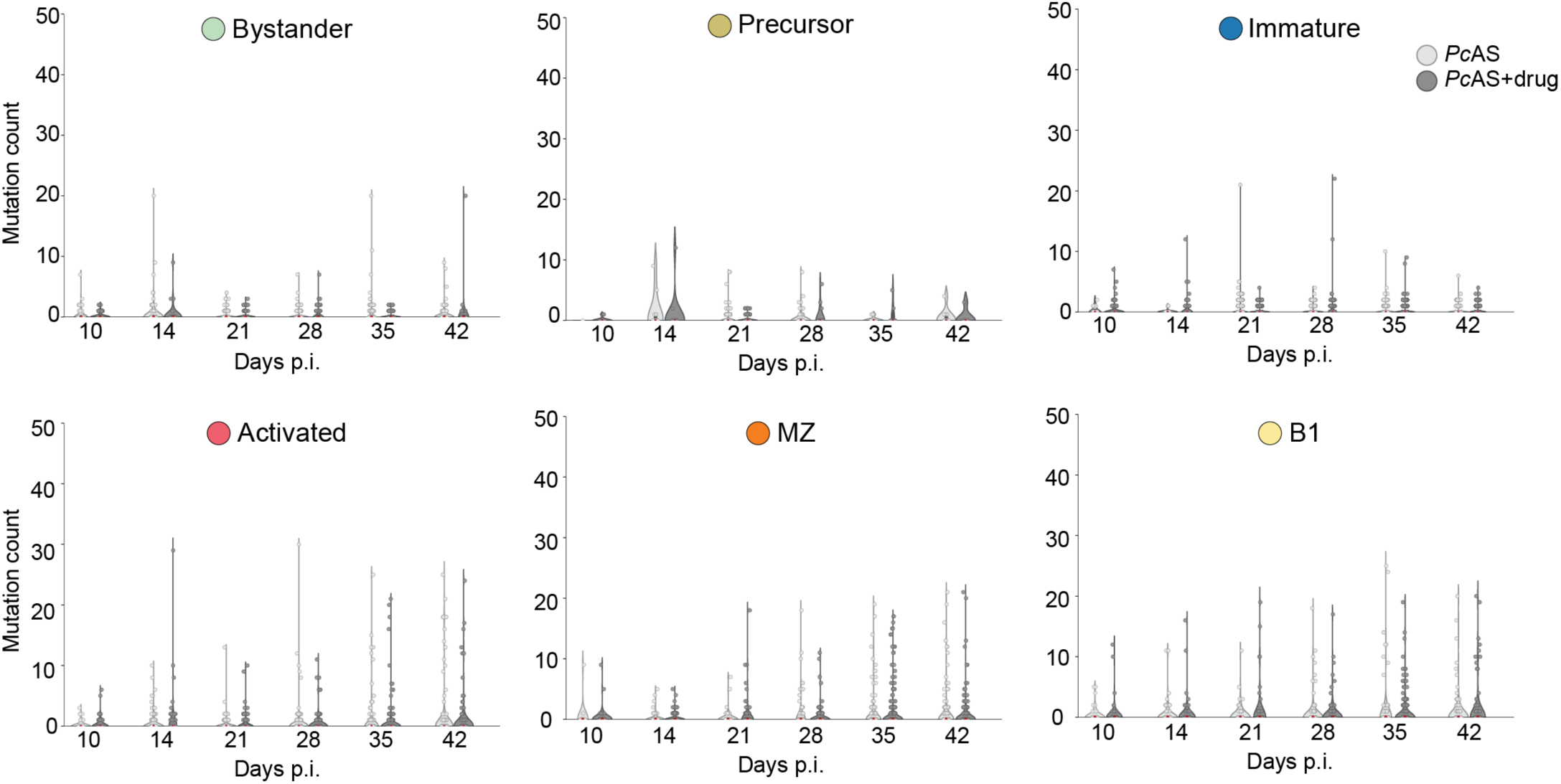
Low BCR mutation counts across multiple B cell populations. Somatic hypermutation analysis of various B cells (naive, precursor, immature, activated, MZ, B1 and bystander B cells) over the course of *Plasmodium* infection, split by treatment. Median number of mutations highlighted in red.

**Extended Data Figure 9:**
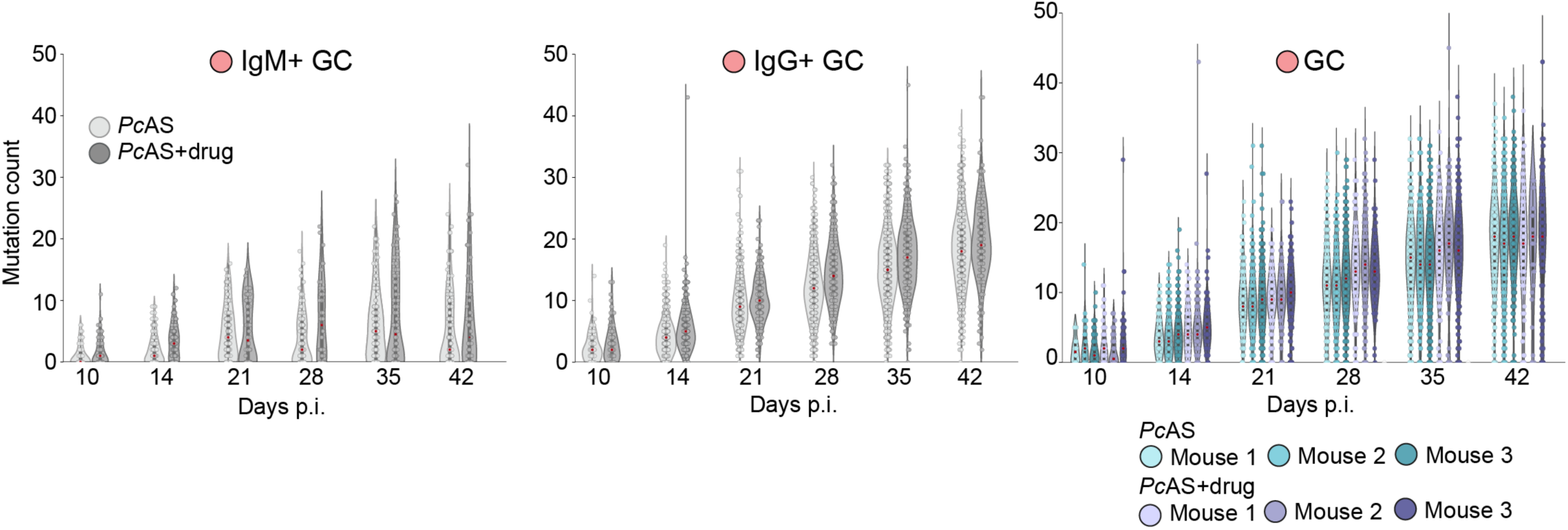
BCR mutation counts in GC B cells over time, as depicted for different isotypes and across individual mice. Somatic hypermutation analysis of GC B cells over the course of *Plasmodium* infection, split by treatment, shown as unswitched (IgM^+^) and switched (IgG^+^) and also shown mutations in all GC B cells split by individual mice. Median number of mutations highlighted in red.

**Extended Data Figure 10:**
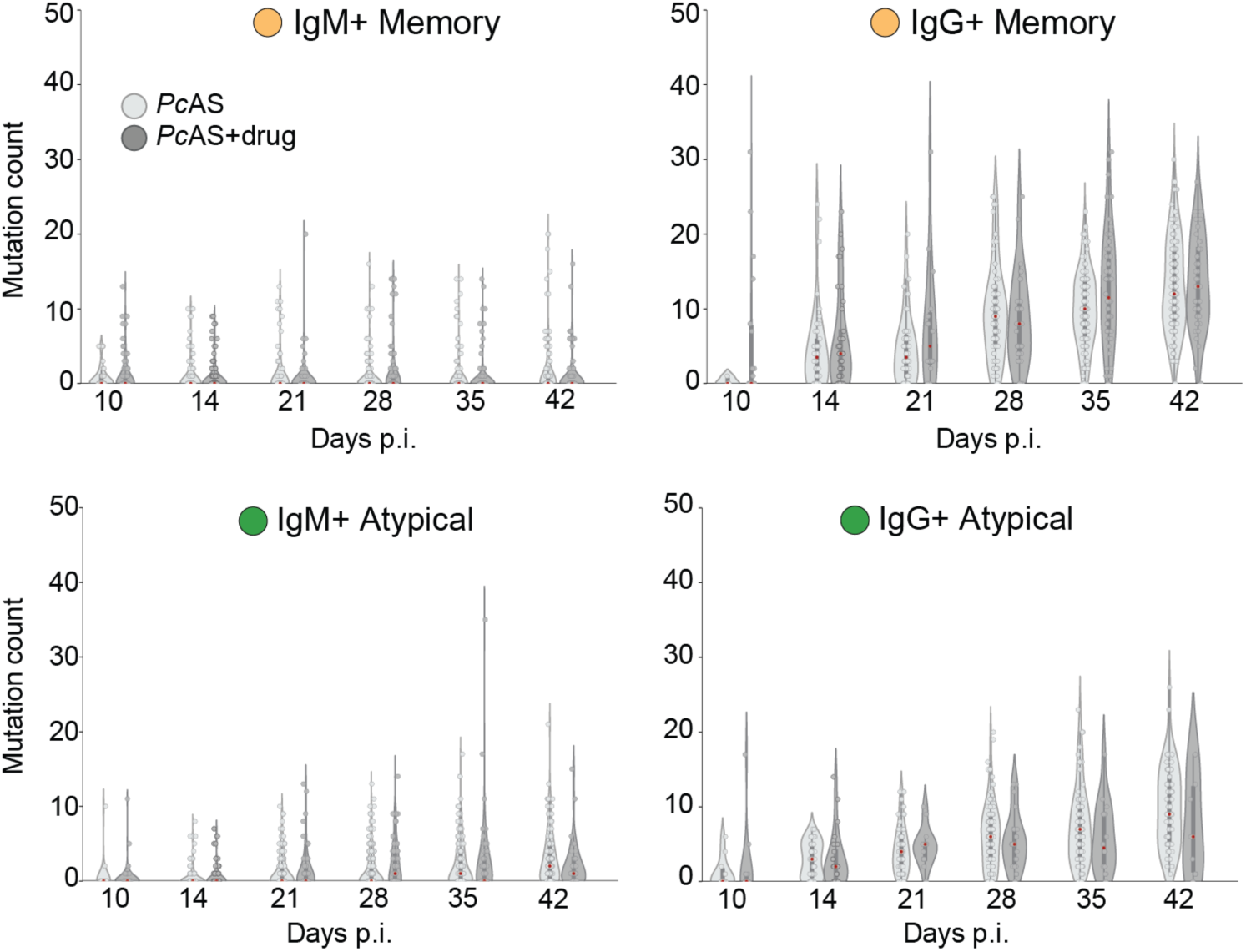
BCR mutations over time for atypical and memory B cells. Somatic hypermutation analysis of memory and atypical B cells split by unswitched (IgM^+^) and switched (IgG^+^) over the course of *Plasmodium* infection, split by treatment. Median number of mutations highlighted in red.

**Extended Data Figure 11:**
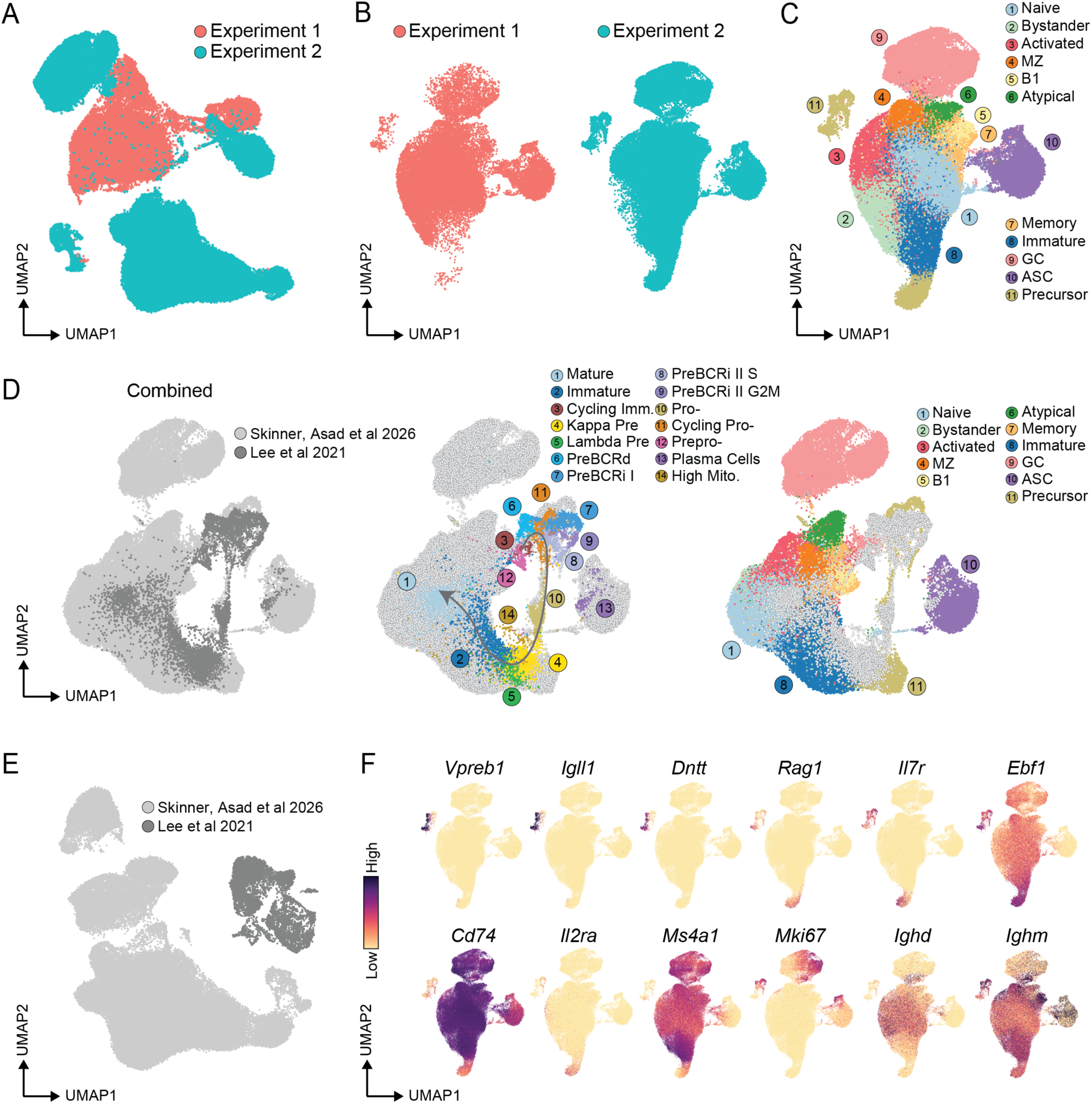
Computational integration with bone marrow scRNAseq data facilitates identification of developing B cell subsets in the spleen. (**A**) scVI integration of experiment 1 and 2 without batch correction. (**B**) Integrated scVI UMAP of experiment 1 and 2, using batch correction, split by experiment. (**C**) Annotated, scVI integrated UMAP of both experimental datasets. (**D**) scVI integration of our day 10-42 B cell dataset with publicly available scRNAseq bone marrow data combined and split by each dataset, with arrow indicating inferred trajectory of Lee et al., 2021 dataset. (**E**) scVI integration of our day 10-42 B cell dataset and publicly available bone marrow B cell scRNAseq dataset, without batch correction. (**F**) Expression of canonical markers of pro-B cells (*Vpreb1, Igll1, Dntt*), cycling pro-B cells (*Mki67*), pre-B cells (*Il2ra*), both pro- and pre-B cells (*Rag1, Il7r, Ebf1, Cd74^lo^, Ighd^-^*) and immature B cells (*Ms4a1, Ighd^-^, Ighm^+^*).

**Extended Data Figure 12:**
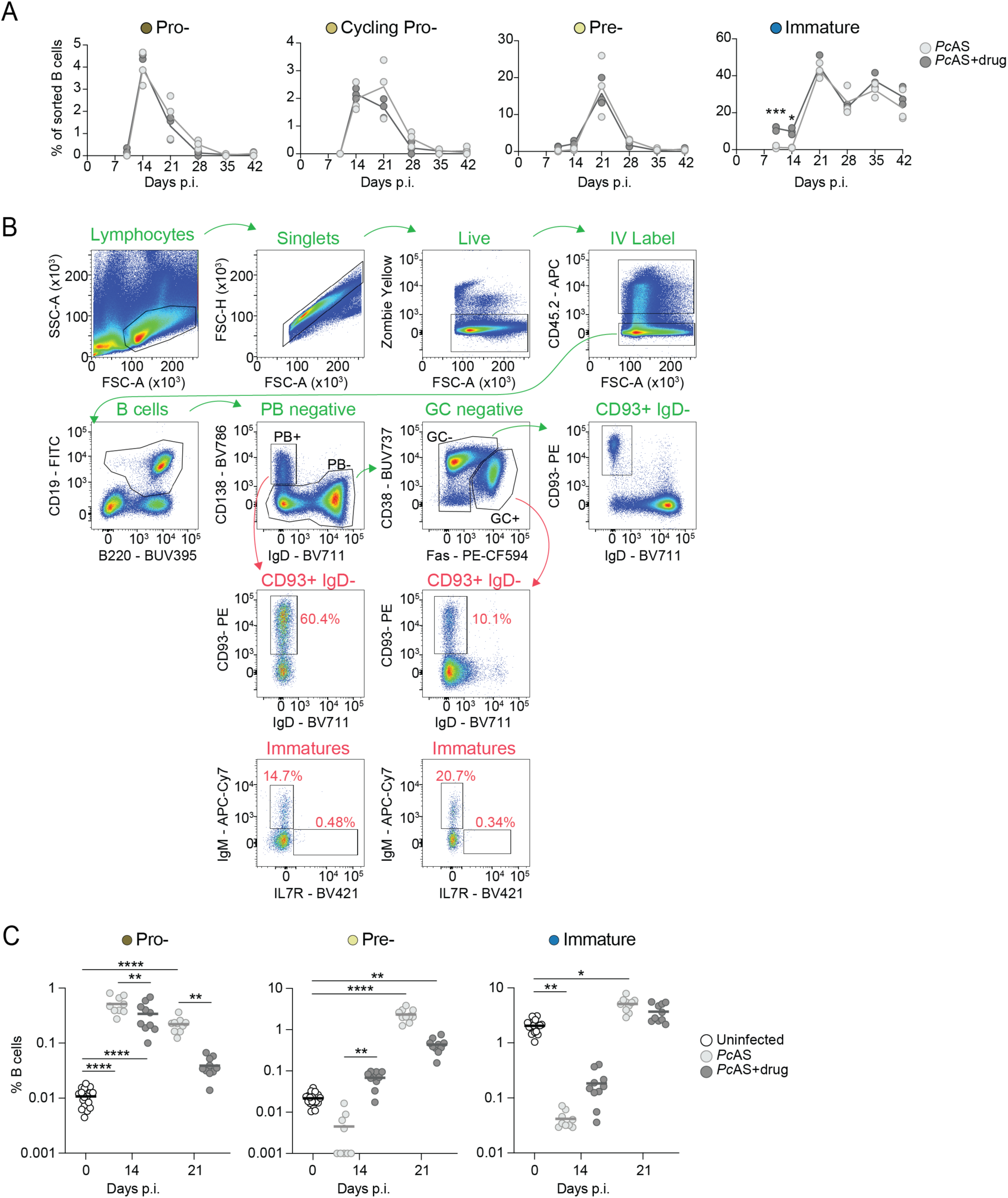
Splenic B cell development during experimental malaria is unaffected by anti-malarial drug treatment. (**A**) Quantification of pro-, cycling pro-, pre- and immature B cells in replicate mice from the integrated UMAP. (**B**) Gating strategy for identifying splenic precursor B cells after *Plasmodium* infection by gating on live CD45.2^-^ CD19^+^ B220^+^ B cells, CD138^-^ IgD^+^ non-plasmablasts, CD38^+^ Fas^-^ non-GCs and IgD^-^ CD93^+^ to look for immature B cells (IgM^+^ IL7R^-^) and pre- and pro-B cells (IgM^-^ IL7R^+^). (**C**) Quantification of flow cytometric analysis of pro-, pre- and immature B cells at 14 and 21 dpi with or without anti-malarial drug treatment. Lines in panel A connect the mean values at each timepoint and analysed by Welch’s *t*-test. Data in panel C is the mean of n=9-20 mice pooled form 2 independent experiments and analysed by one-way ANOVA with Tukey’s multiple comparisons test (pro-B cells) or Kruskal-Wallis test with Dunn’s multiple comparisons test (pre- and immature B cells). \**P* <0.05, \*\**P* <0.01, \*\*\**P* <0.001, \*\*\*\**P* <0.0001.

**Extended Data Figure 13:**
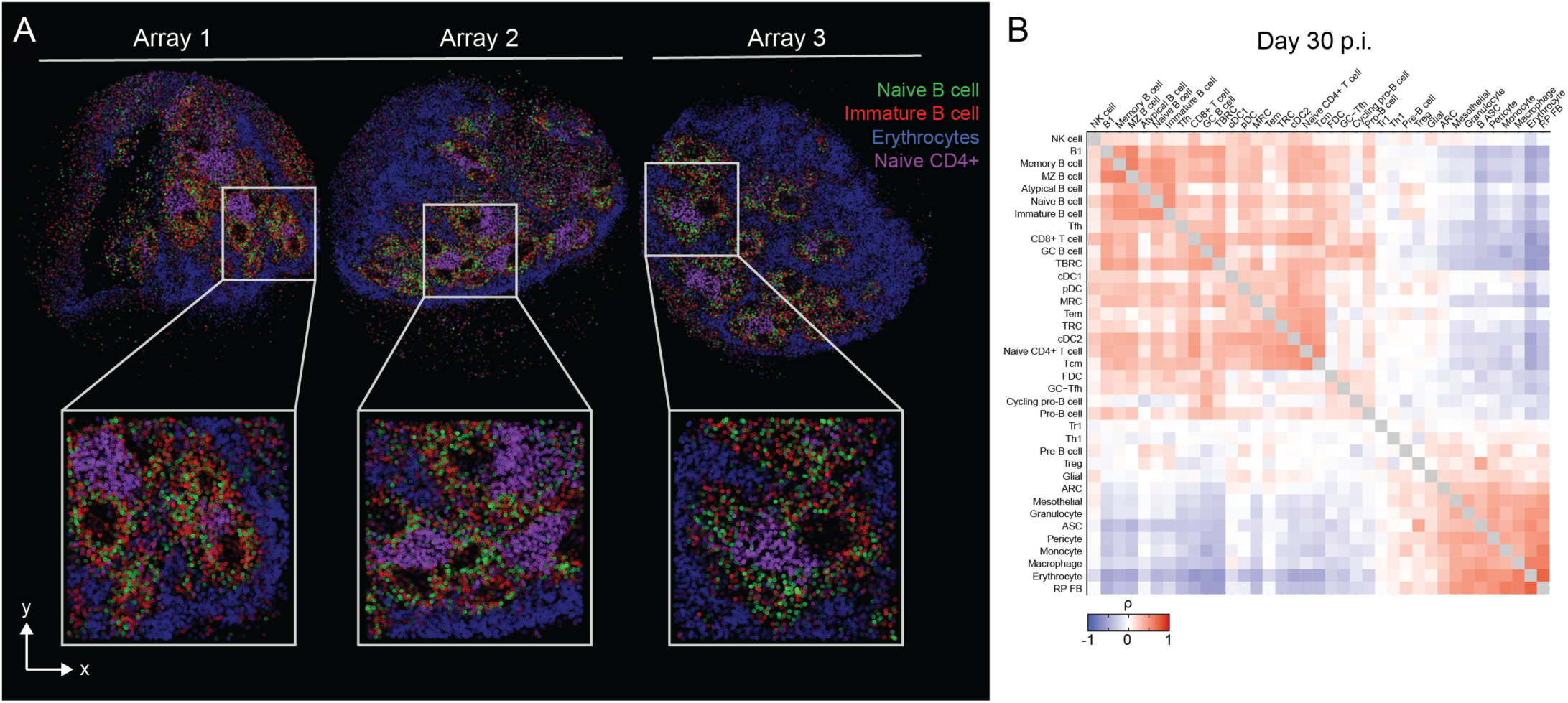
Multiple spatial transcriptomic arrays at 30 dpi reveal co-localisation of immature B cells with naïve B cell follicles in the spleen. (**A**) RCTD spatial deconvolution in 30 dpi spleens and three independent spleen sections. Arrays 1 and 2 are from the same mouse spleen. Array 3 is from same spleen as in Fig. 6C. (**B**) Spearman’s correlation analysis of all cell co-localisations from all 4 arrays.

**Extended Data Figure 14:**
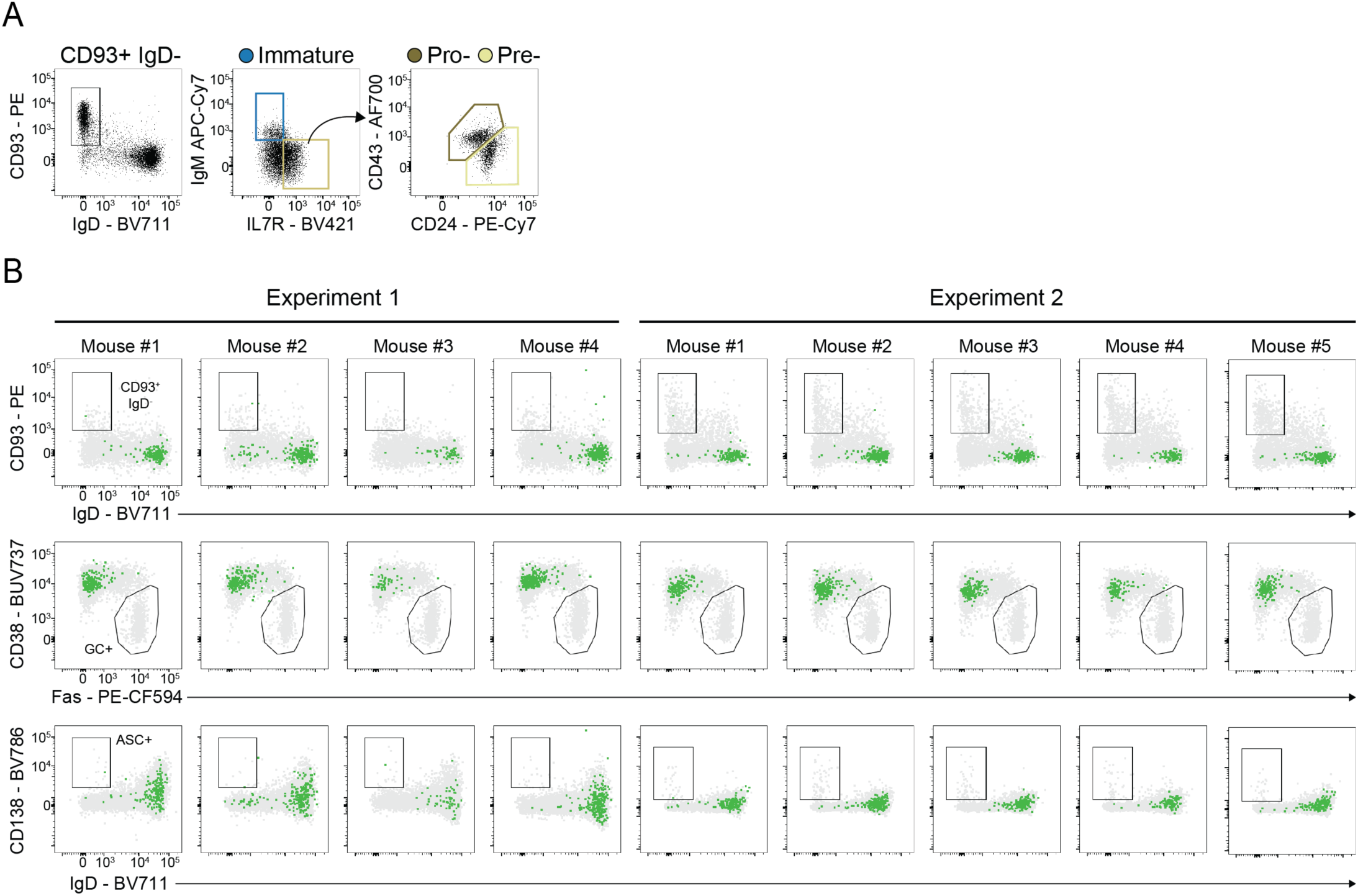
QC and summary data illustrating splenic precursor B cells develop into mature B cells *in vivo*. (**A**) Representative flow cytometry panel of splenocytes gated on CD19^+^ B220^+^ CD138^-^ CD38^+^ Fas^-^for CD93^+^ IgD^-^ B cells, followed by immature B cells (IgM^+^ IL7R^-^), pro-B cells (IgM^-^ IL7R^+^, CD43^hi^ CD24^int^) and pre-B cells (IgM^-^ IL7R^+^, CD43^int^ CD24^hi^) in mice at 21 dpi with *Py*17XNL. (**B**) Flow cytometry panels of CD93^+^ IgD^-^ B cells, GC B cells and ASCs at 21 days post-transfer of GFP^+^ CD93^+^ IgD^-^ splenic precursor B cells (green cells). Cells in grey are endogenous cells from host mice at day 42 post *Py*17XNL infection.

## Supplementary Method Figures

**Supplementary Method Figure 1:**
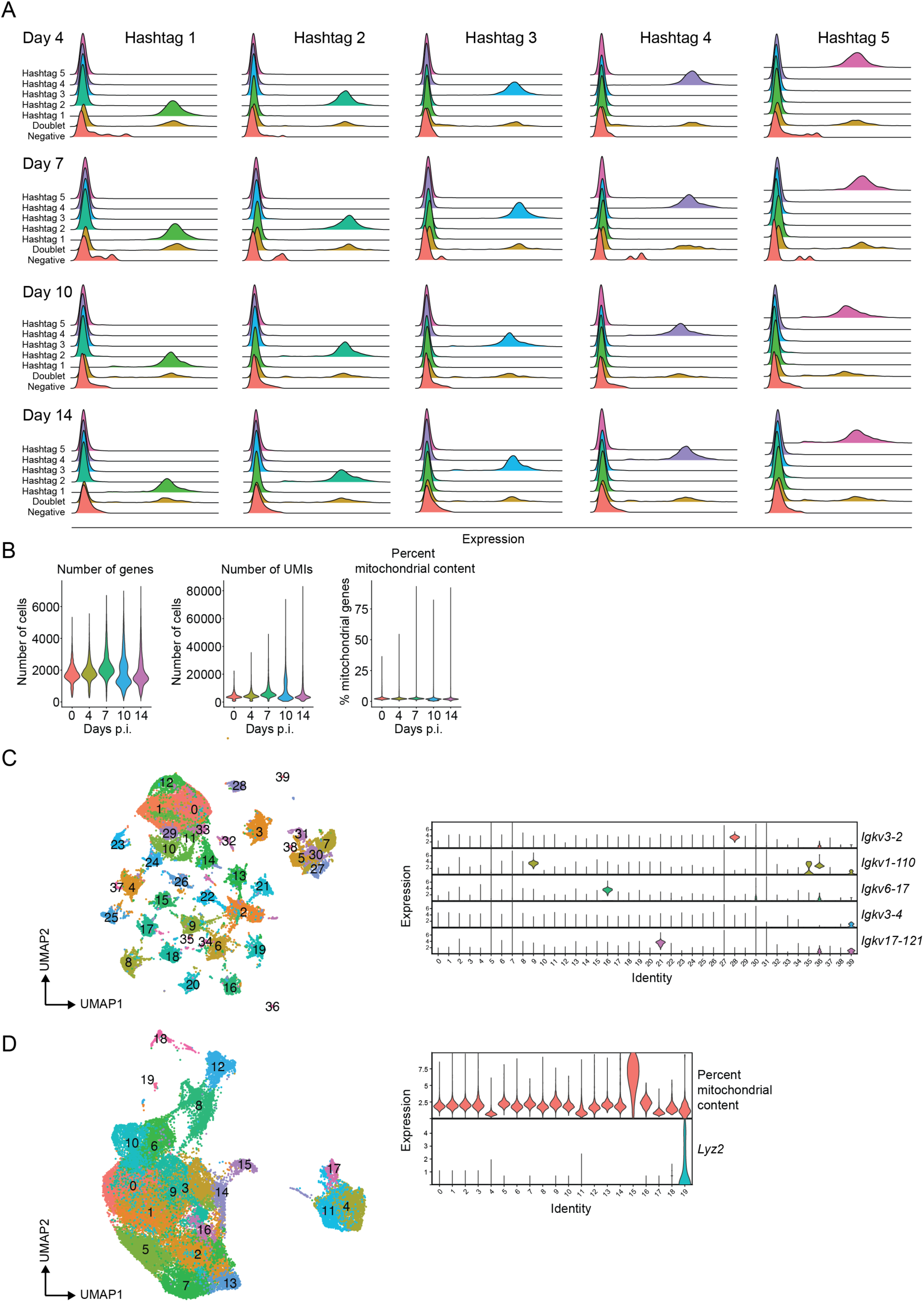
Quality control for B cell scRNAseq Experiment 1: 0-14 dpi. (**A**) Ridge plots for each hashtag showing expression of hashtag oligos for singlets (hashtags 1-5), doublets and negative cells for each timepoint. Note, hashtag 1 in the day 4 group is the day 0, naive B cell sample. (**B**) Violin plots of number of genes, number of UMIs and percent mitochondrial content prior to pre-processing of data. (**C**) UMAP of entire B cell dataset after high-dimensional reduction and clustering prior to removal of BCR-related genes including violin plots showing distribution of expression of a random selection of BCR related genes for each cluster. (**D**) UMAP of entire B cell dataset after removal of BCR-related genes and after high-dimensional reduction and clustering. Violin plots show the distribution of percent mitochondrial content and *Lyz2* (monocytes) expression per cluster.

**Supplementary Method Figure 2:**
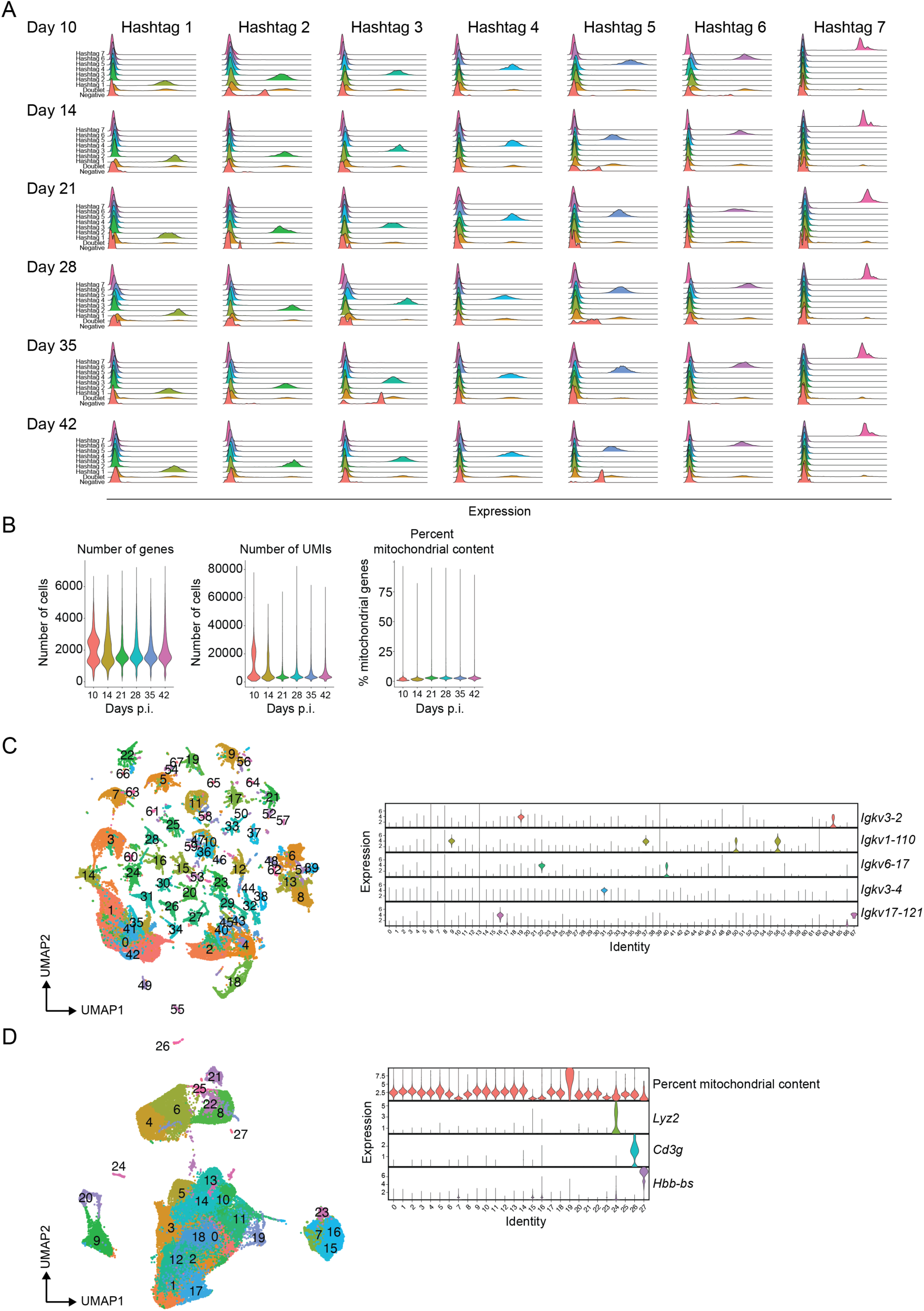
Quality control for B cell scRNAseq Experiment 2: 10-42 dpi. (**A**) Ridge plots for each hashtag showing expression of hashtag oligos for singlets (hashtags 1-7), doublets and negative cells for each timepoint. (**B**) Violin plots of number of genes, number of UMIs and percent mitochondrial content prior to pre-processing of data. (**C**) UMAP of entire B cell dataset after high-dimensional reduction and clustering prior to removal of BCR-related genes including violin plots showing distribution of expression of a random selection of BCR related genes for each cluster. (**D**) UMAP of entire B cell dataset after removal of BCR-related genes and after high-dimensional reduction and clustering. Violin plots show the distribution of percent mitochondrial content and *Lyz2* (monocytes), *Cd3g* (T cells) and *Hbb-bs* (erythrocytes) expression per cluster.

**Supplementary Method Figure 3:**
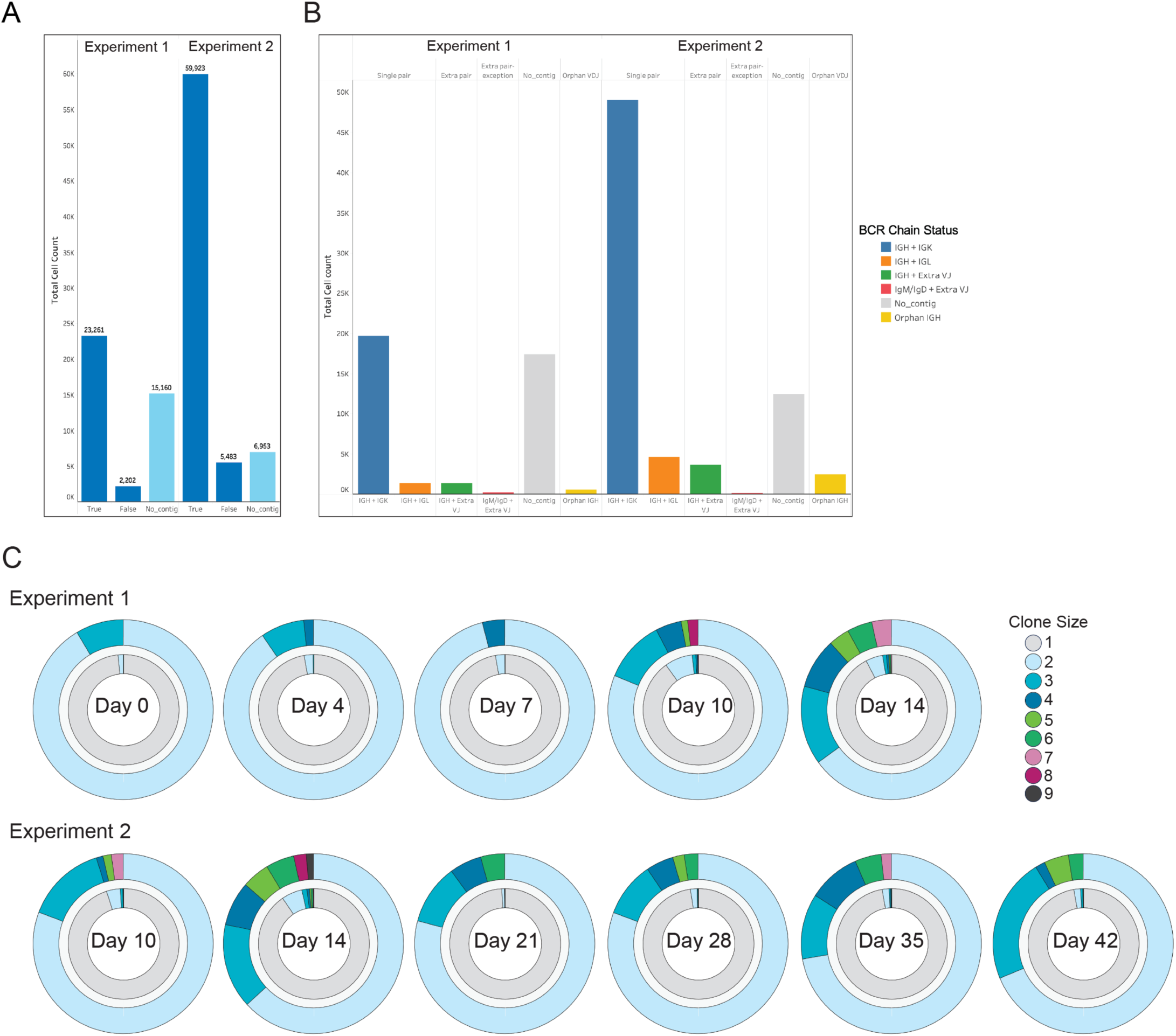
Quality control for clonal analysis from single-cell BCRseq. (**A**) Bar graph showing distribution of cells which passed Dandelion quality control pipeline for each experiment. (**B**) Bar graph showing distribution of BCR chains in each experiment. (**C**) Donut plots showing all clones including unexpanded and expanded clones (inner donut) and expanded clones only (outer donut) at each timepoint for each experiment.

